# A RALF-Brassinosteroid morpho-signaling circuit regulates *Arabidopsis* hypocotyl cell shape

**DOI:** 10.1101/2024.03.28.587199

**Authors:** David Biermann, Michelle von Arx, Kaltra Xhelilaj, David Séré, Martin Stegmann, Sebastian Wolf, Cyril Zipfel, Julien Gronnier

## Abstract

Plant cells survey and modulate their cell wall to control their shape and anisotropic growth. Signaling mediated by the plant steroid hormones brassinosteroids (BR) plays a central role in coordinating cell wall status and cell growth, and alterations in the cell wall – BR feedback loop leads to life-threatening defects in tissue and cellular integrity. How the status of the cell wall is relayed to BR signaling remains largely unclear. Increasing evidence shows that RAPID ALKALANIZATION FACTORs (RALFs), a class of secreted peptides, play structural and signaling roles at the cell surface. Here we show that perception of RALF23 promotes the formation and signaling of the main BR receptor complex formed by BRASSINOSTEROID INSENSITIVE 1 (BRI1) and BRI1-BRASSINOSTEROID INSENSITIVE1-BRASSINOSTEROID-ASSOCIATED KINASE 1

(BAK1). The loss of the plasma membrane-localized RALF receptor complex FERONIA (FER)-LORELEI LIKE GPI-anchor protein 1 (LLG1) leads to defects in cell expansion and anisotropy, as well as uncontrolled BRI1-BAK1 complex formation and signaling. RALF23 bioactivity relies on pectin status and its perception induces changes in pectin composition and the activity of pectin-modifying enzymes. Our observations suggest a model in which RALF23 functions as a cell wall-informed signaling cue initiating a feedback loop that solicits BR signaling, modifies the cell wall, and coordinates cell morphogenesis.

**Highlights:** -The RALF receptor complex FER-LLG1 regulates cell anisotropic growth

-RALF23 promotes BRI1-BAK1 complex formation and signaling

-RALF23 functions as a cell wall-informed and wall-modifying signaling cue

## Introduction

Plant cells are encased by the cell wall, a rigid polysaccharide-rich matrix providing tensile strength and controlling cell shape and integrity (Anderson and Kieber 2020). During development, expanding cells face a life-or-death dilemma: they must precisely control the modifications of the cell wall to allow anisotropic growth while preserving cell wall integrity (Vaahtera et al. 2019). Accumulating evidence indicates that this tight balance is controlled by wall-derived cues that are perceived as informative signals at the cell wall–plasma membrane interface to initiate cell wall signaling (CWS) (Hématy et al. 2007; Mecchia et al. 2017; Van der Does et al. 2017; Engelsdorf et al. 2018; Herger et al. 2019; Chaudhary et al. 2020; Wolf 2022; Liu et al. 2023). Among the recently identified CWS components, the RAPID ALKALINIZATION FACTOR (RALF) family of secreted peptides and its binding partners play a pivotal role in regulating growth (Srivastava et al. 2009; Gonneau et al. 2018; Dünser et al. 2019; Li et al. 2022; Gupta et al. 2024), reproduction (Ge et al. 2017; Mecchia et al. 2017; Zhong et al. 2022; Lan et al. 2023; Zhou et al. 2023), responses to abiotic stresses (Feng et al. 2018; Zhao et al. 2018, 2021) and immunity (Guo et al.; Stegmann et al. 2017; Song et al. 2021; Gronnier et al. 2022; Tang et al. 2022). Recent studies show that positively charged RALF peptides associate with cell wall carbohydrates in the form of de-esterified pectin (Moussu et al. 2020, 2023; Schoenaers et al. 2023; Liu et al. 2024; Rößling et al. 2024). RALF-pectin mers associate with cell wall-bound LEUCINE-RICH REPEAT EXTENSIN (LRX) proteins (Mecchia et al. 2017; Dünser et al. 2019; Herger et al. 2020; Moussu et al. 2020, 2023) and plasma membrane-localized complexes composed of *Catharanthus roseus* RECEPTOR-LIKE KINASE 1-LIKE (CrRLK1L) and LORELEI-LIKE glycosylphosphatidylinositol-anchored (LLG) proteins (Haruta et al. 2014; Li et al. 2015; Gonneau et al. 2018; Ge et al. 2019; Xiao et al. 2019; Liu et al. 2024). In the cell wall of tip-growing cells, RALF-pectin-LRX complexes assemble into a reticulated network providing structural support for cellular expansion (Moussu et al. 2023; Schoenaers et al. 2023). At the cell surface, RALF perception has been linked to the initiation of several signaling events such as changes in apoplastic pH (Pearce et al. 2001; Morato do Canto et al. 2014; Li et al. 2022), the production of reactive oxygen species (Stegmann et al. 2017; Abarca et al. 2021), and an influx of calcium (Haruta et al. 2014; Gjetting et al. 2020; Gao et al. 2023). We previously reported that perception of RALF23 by the plasma membrane-localized receptors LLG1 and the CrRLK1L FERONIA inhibits complex formation of the immune LEUCINE RICH REPEAT (LRR) receptor kinase (RK) FLAGELLIN SENSING 2 (FLS2) and its co-receptor BRASSINOSTEROID INSENSITIVE 1 (BRI1)-ASSOCIATED KINASE 1 (BAK1, also known as SOMATIC EMBRYOGENESIS RECEPTOR KINASE 3, SERK3) (Stegmann et al. 2017; Xiao et al. 2019) and that such inhibitory function is linked to the regulation of FLS2 and BAK1 plasma membrane organization (Gronnier et al. 2022). It has recently been shown that this effect is not limited to FLS2 and BAK1 and that perception of RALF23 induces global changes in membrane organization (Yu et al. 2020; Chen et al. 2023b; Smokvarska et al. 2023; Liu et al. 2024); notably impacting the organization of additional cell surface receptors, such as the LRR-RK BRI1 (Liu et al. 2024). While the effect of RALF perception on immune signaling is well characterized (Stegmann et al. 2017; Song et al. 2018; Gronnier et al. 2022; Tang et al. 2022; Chen et al. 2023a), the functional consequences of RALF perception for other RK signaling pathways and potential link with CWS remain however unexplored. BRI1, the main receptor for the plant steroid hormones brassinosteroids (BR), forms a ligand-induced complex with SERKs such as BAK1/SERK3 (Santiago et al. 2013; Sun et al. 2013; Ma et al. 2016). BRI1-BAK1 complex formation triggers a signaling cascade that involves the de-phosphorylation of the transcription factors of the BRI1 EMS-SUPRESSOR 1 (BES1) and BRASSINAZOLE RESISTANT 1 (BZR1) family (Wang et al. 2002; Yin et al. 2002), and initiates a transcriptional reprogramming regulating growth (Kim and Wang 2010; Belkhadir and Jaillais 2015; Planas-Riverola et al. 2019; Nolan et al. 2022). It is becoming clear that BR signaling functions at the crossroad between CWS and growth (Wolf 2022). Indeed, alteration of pectin methylation status triggers BR signaling (Wolf et al. 2012, 2014), BR signaling modulates the expression of a multitude of cell wall-associated transcripts (Sun et al. 2010; Yu et al. 2011), as well as the activity of wall modifying enzymes (Qu et al. 2011; Xie et al. 2011; Sánchez-Rodríguez et al. 2017), and the inhibition of BR signaling in a mutant affecting pectin modification leads to life-threatening defects in tissue and cellular integrity (Kelly-Bellow et al. 2023). Despite the apparent vital importance of the interplay between the cell wall and BR signaling, little is known in regard to the underpinning molecular actors. In recent years, however, several lines of evidence point towards the predominant role of wall monitoring cell-surface receptors in this process. Indeed, the RECEPTOR-LIKE PROTEIN 44 (RLP44) and the WALL-ASSOCIATED KINASE (WAK)-like RLK RESISTANCE TO FUSARIUM OXYSPORUM 1 (RFO1)/WAKL22 mediate the activation of BRI1 signaling in response to pectin modification (Wolf et al. 2014; Huerta et al. 2023). In rice (*Oryza sativa*) changes in pectin alleviate the inhibition of OsBRI1 signaling by the WALL-ASSOCIATED KINASE 11 (OsWAK11) (Yue et al. 2022). Here we show that RALF23 promotes BRI1-BAK1 complex formation and signaling. We observed that the loss of *FER* and *LLG1* leads to uncontrolled BRI1-BAK1 complex formation and signaling, as well as to defects in cell expansion and anisotropy. RALF23 bioactivity relies on pectin status and its perception induces changes in pectin composition and the activity of pectin-modifying enzymes. Our observations suggest a model in which RALF23 functions as a cell-wall informed signaling cue, initiating a feedback loop that solicitates BR signaling, modifies the cell wall, and coordinates cell morphogenesis.

## Results and discussion

### FER inhibits BRI1-BAK1 complex formation and signaling

The *FER* loss of function mutant *fer-2* was reported to be hypersensitive to exogenous treatment with 24-epi-brassinolide (BL), a potent brassinosteroid (Deslauriers and Larsen 2010). To confirm these observations, we used an additional knock-out allele for *FER*, *fer-4*, and a corresponding complementation line expressing FER-GFP under the control of its native promotor (Duan et al. 2010). In hypocotyl elongation assays, we observed that *fer-4* is hypersensitive to BL treatment and that expression of FER-GFP restored BL responsiveness to wild-type levels (Figure 1A). Since FER regulates ligand-induced complex formation between FLS2 and BAK1 (Stegmann et al. 2017), we asked whether the increased BL responsiveness observed in *fer-4* can be explained by changes in BRI1-BAK1 complex formation. In co-immunoprecipitation experiments, we observed a constitutive increase in the association between BRI1-GFP and BAK1 in *fer-4* compared to WT (Figure 1B). To corroborate these observations, we monitored BES1 phosphorylation status and transcript accumulation of two BR-responsive marker genes negatively regulated by BRs, *CPD* and *DWF4* (Albrecht et al. 2012a). We observed a constitutive decrease in BES1 phosphorylation as well as a constitutive decrease in the transcript accumulation of *CPD* and *DWF4* in *fer-4*, indicative of an increase in BR signaling (Figure 1C-D). Taken together, these results show that FER inhibits BR signaling by regulating BRI1-BAK1 complex formation. Co-immunoprecipitation experiments suggest that FER is not in close proximity to BRI1 (Figure S1) implying that regulation of BRI1-BAK1 association by FER may be indirect.

**Figure 1.**
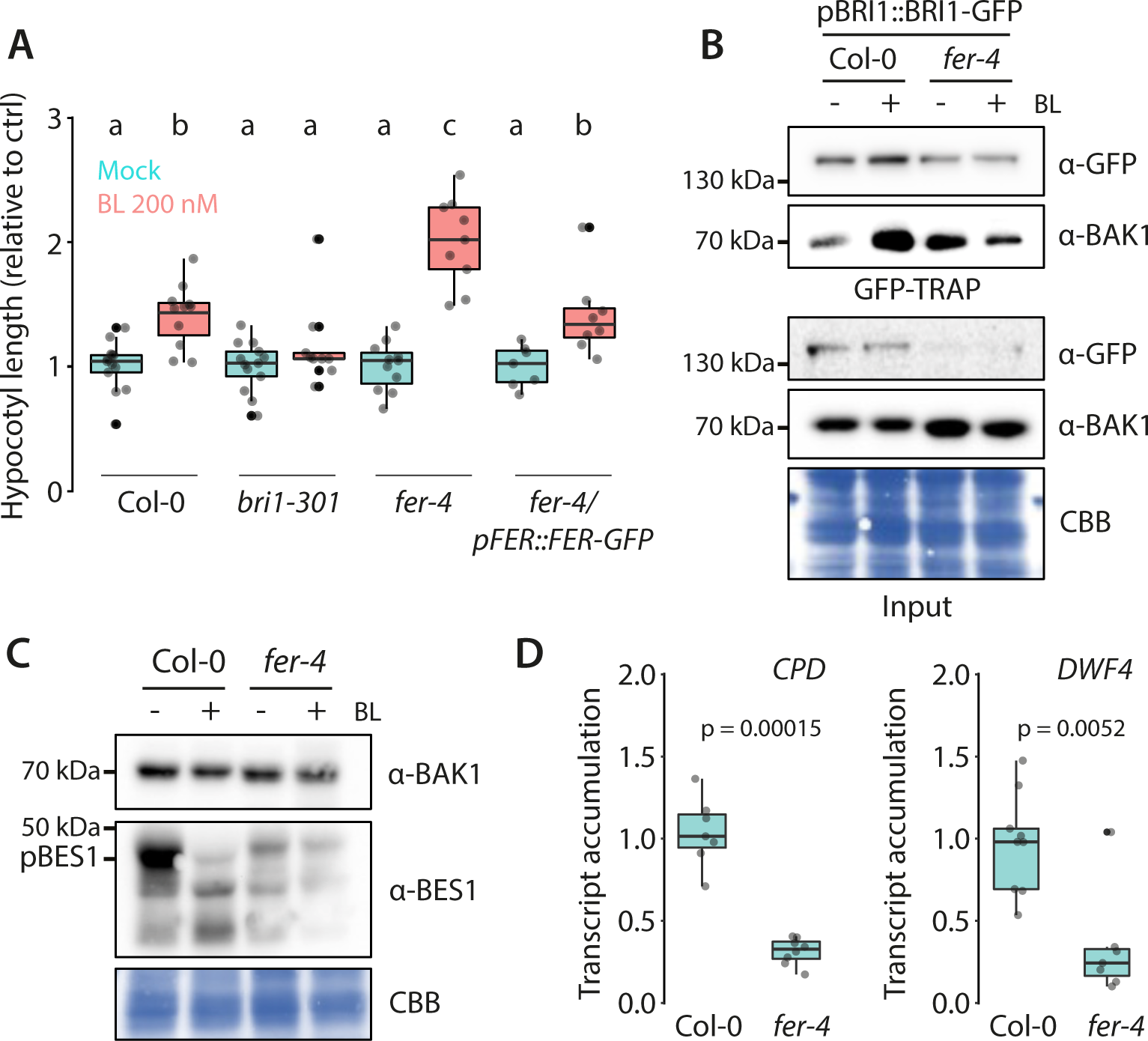
FERONIA inhibits BRI1-BAK1 complex formation and signaling. **A.** Quantification of hypocotyl length of five-day-old Arabidopsis seedlings grown on half MS medium containing 200 nM brassinolide (BL) or corresponding mock control solution (EtOH). Scattered data points indicate measurements of individual seedlings. Conditions that do not share a letter are significantly different in Dunn’s multiple comparison test (p<0.0001). **B.** Immunoprecipitation of BRI1-GFP in Arabidopsis seedlings after treatment with 1 µM BL or corresponding mock control solution (EtOH) for 90 minutes. Membranes were probed with anti-GFP or anti-BAK1 antibodies. Membrane stained with Coomassie brilliant blue (CBB) is presented to show equal loading. Similar observations were made in at least three independent experiments. **C.** BES1 de-phosphorylation 30 min after treatment of twelve-day-old seedlings with 200 nM of BL, as shown by western blot analysis using anti-BES1 antibodies. Membranes were probed with anti-BAK1 antibodies, and subsequently stained with Coomassie brilliant blue (CBB) as loading controls. **D.** Quantitative real-time PCR of *CPD* and *DWF4* transcripts from twelve-day-old seedlings. Scattered data points indicate measurements of individual biological samples, each corresponding to 2-3 seedlings pooled, obtained from three independent experiment. *U-BOX* was used as a house keeping gene and values are expressed relative to Col-0. P-values report Mann-Whitney statistical test.

### FER regulates hypocotyl cell anisotropic growth and the activity of pectin modifying enzymes

FER was shown to contribute to the mechanical integrity in the Arabidopsis root and shoot (Shih et al. 2014; Malivert et al. 2021). We set to investigate the potential effect of loss of *FER* on cell growth and morphology in the hypocotyl of light-grown seedlings. Microscopic observation of *fer-4* revealed defects in cell shape (Figure 2, Figure S2). To gain quantitative information on these morphological defects we performed automated high resolution 3D confocal imaging of optically cleared samples and computed 3D volumes of the hypocotyl epidermis using MorphographX (Barbier de Reuille et al. 2015). Compared to wild-type, *fer-4* showed a pronounced increase in cell volume and a decrease in cell anisotropy (Figure 2 A-C). Moreover, we repeatedly observed cellular outgrowths and loss of cellular integrity in *fer-4*, both in fixed and living samples (Figure S2). We conclude that *FER* is genetically required to control anisotropic growth of hypocotyl epidermal cells. In the cell wall, pectin forms a complex and dynamic meshwork that actively participates in shaping plant cells, although the exact contribution of pectin to morphogenesis remains enigmatic (Bidhendi et al. 2019; Majda et al. 2019; Cosgrove and Anderson 2020; Haas et al. 2020). Changes in pectin esterification degree correlates with wall biophysical properties (Peaucelle et al. 2011, 2015) and the overexpression of pectin modifying enzymes, pectin methyl esterases (PME) or PME inhibitors (PMEI) is sufficient to alter cell morphology (Wolf and Greiner 2012; Daher et al. 2018). Loss of *FER* is associated with a decrease in de-esterified pectin at the filiform apparatus of synergids cells (Duan et al. 2020). We wondered whether defects in cell shape of the hypocotyl epidermis observed in *fer-4* are linked to changes in pectin status in vegetative tissue. We used a dot-blot assay, probing isolated pectin with LM20, JIM5 and LM19 antibodies which present distinct affinity towards highly esterified, partially esterified, and mostly de-esterified pectin, respectively (Christiaens et al. 2011). We observed a decrease in signal of all antibodies used to probe *fer-4* cell wall fractions, indicating that loss of *FER* alters pectin methylation status (Figure 2C). To corroborate these observations, we examined the activity of PMEs, the enzymes responsible for pectin de-esterification (Wolf and Greiner 2012). We observed an increase in PME activity in *fer-4* proteinaceous cell wall extracts (Figure 2D). Altogether, these observations indicate that FER is genetically required to control wall modifications and cell anisotropy. Since BR is a major determinant promoting cell growth and expansion (Wang et al. 2012; Fridman and Savaldi-Goldstein 2013), we asked whether increased BR signaling causes the increase in cell volume and the loss of anisotropy observed in *fer-4*. We crossed *fer-4* with *bak1-4*, a *BAK1* knock-out allele (Chinchilla et al. 2007) and *bri1-301*, an hypomorphic allele of *BRI1* (Xu et al. 2008) (Figure 3A). As expected, *bri1-301* and *bak1-4* were hyporesponsive to BL treatment (Figure 3B). We observed that introducing *bri1-301* or *bak1-4* alleles in *fer-4* alleviates *fer-4* hypersensitivity to exogenous BL treatment (Figure 3B) indicating that they effectively mitigate the increase in BR signaling observed in *fer-4*. However, quantitative analysis of hypocotyl epidermal cell 3D morphology showed that introducing *bak1-4* or *bri1-301* in *fer-4* does not alleviate cell volume and cell shape defects of *fer-4* (Figure 3C-D). In contrast, cell anisotropy appears more severely affected in *fer-4/bak1-4* and *fer-4/bri1-301* than in *fer-4* (Figure 3C-D). These results indicate that increased BR signaling is not causal for the *fer-4* morphological defects. On the contrary, these observations suggest that the increase in BR signaling could rather attenuate the defects caused by the loss of *FER*. Corroborating this hypothesis, we observed that BL treatment alleviated fer-4 defects in cell anisotropy (Figure S3). Altogether, these observations suggest a tight link between FER, cell morphology and BR signaling.

**Figure 2.**
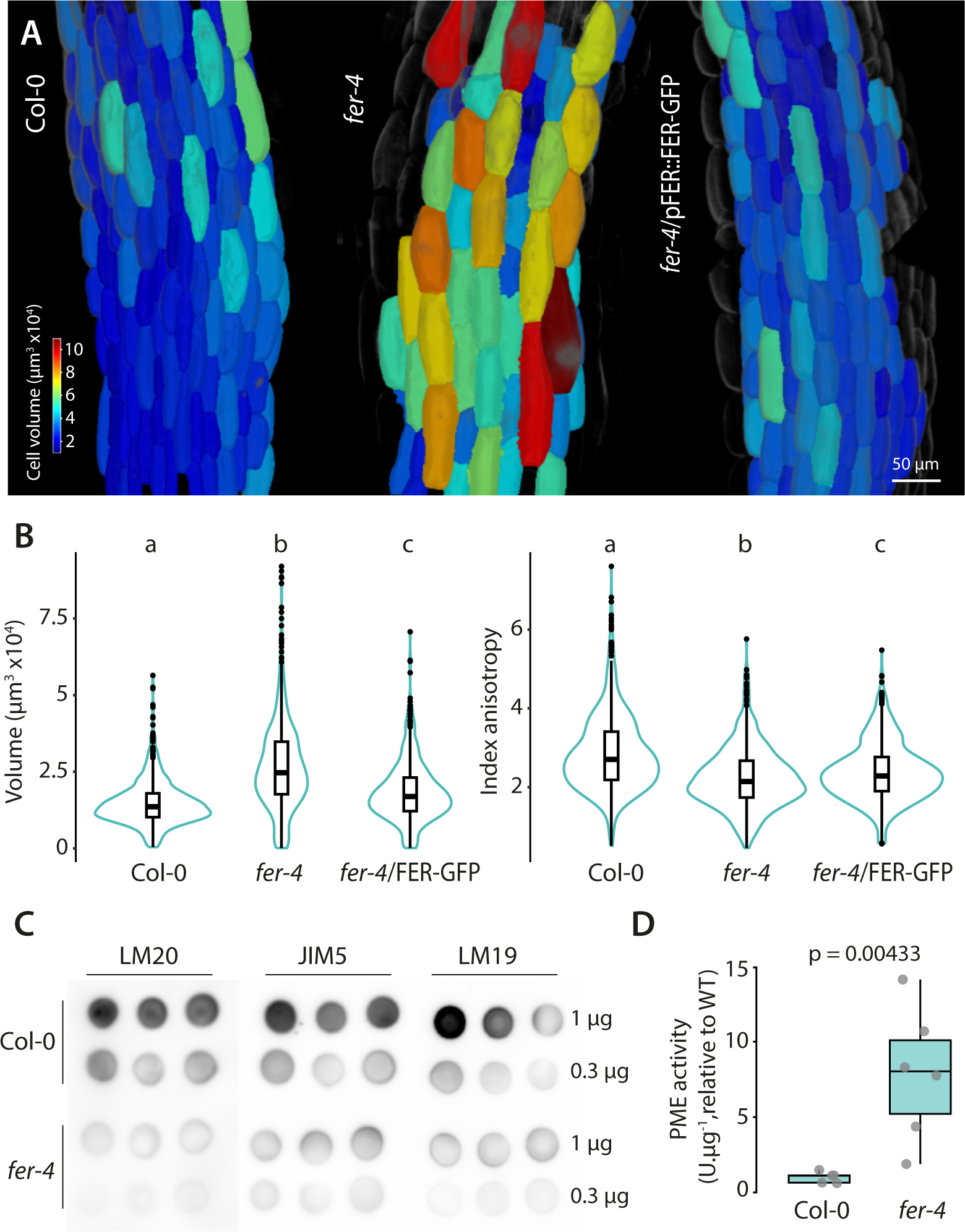
FERONIA regulates cell anisotropic growth and pectin methylation status. **A.** 3D segmentation of the hypocotyl epidermal cells of fixed five-days-old seedlings stained with SR2200. Cells are colored according to their volume. **B.** Quantification of cell volume and anisotropy in Col-0 and *fer-4*. Graphs are combined violin and box plots, n= 647 – 760 cells from 5-6 seedlings. Individual scatter points show outliers. Conditions which do not share a letter are significantly different according to one-way Anova with Tukeys post-hoc HSD test (p < 0.05). **C.** Dot-blot showing abundance of esterified (LM20), partially de-esterified (JIM5) and mostly de-esterified (LM19) pectin in Col-0 and *fer-4*; µg indicates the quantity of total sugars spotted on membrane. 3 independent biological replicates are shown. **D.** Relative PME activity determined by gel diffusion assay. Scatter points show 3 independent biological replicates from 2 experiments, p-value was determined by pairwise Mann-Whitney test.

**Figure 3.**
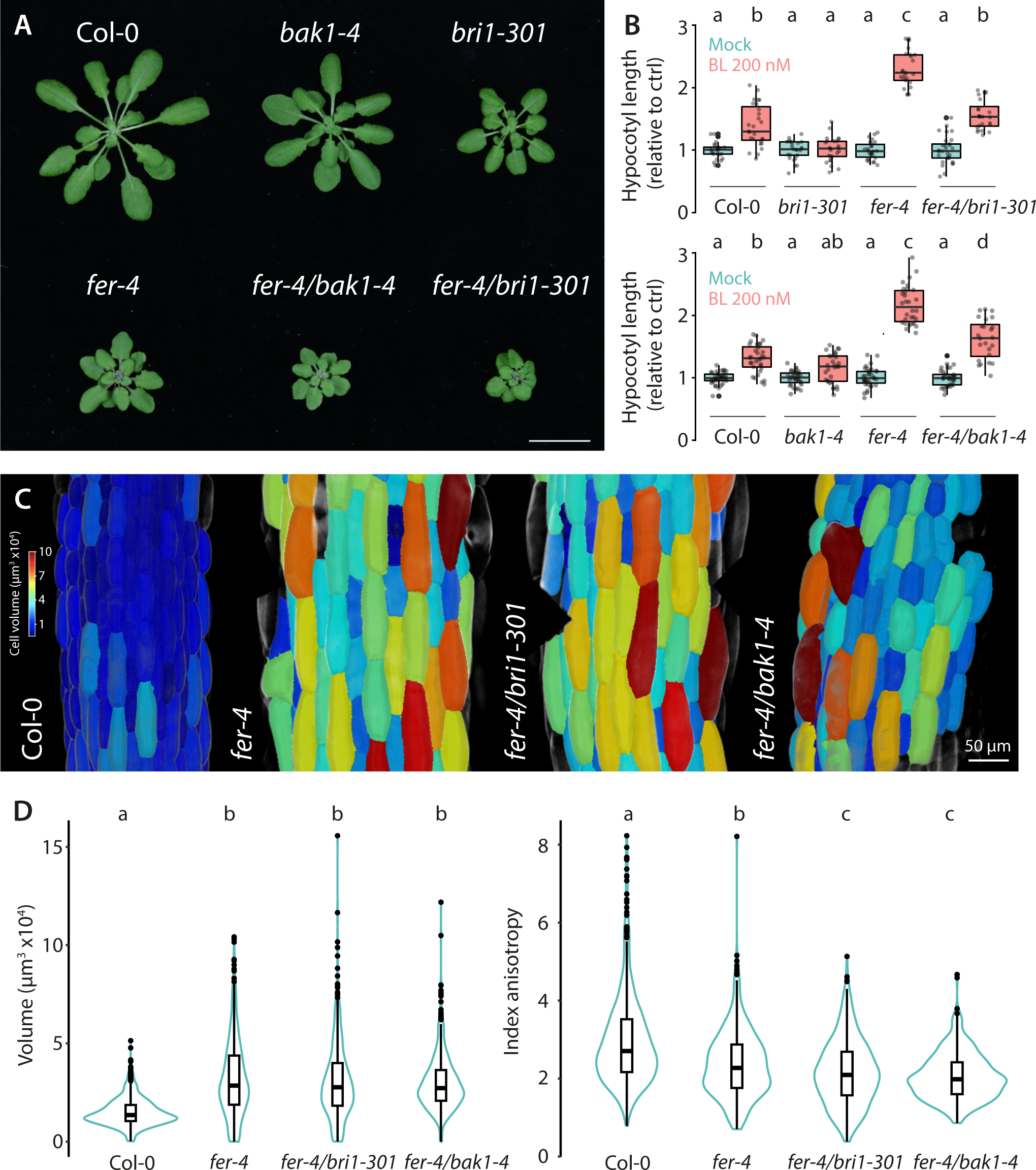
Increase in BR signaling does not underly morphological defects of *fer-4*. **A.** Rosettes of four-week-old Arabidopsis plants grown in a 16-hour light regime. **B.** Quantification of hypocotyl lengths of five-day-old seedlings grown on half MS medium containing 200 nM brassinolide (BL) or corresponding mock control solution (EtOH). Graphs are boxplots, scattered data points show individual measurements pooled from three independent experiments. Conditions that do not share a letter are significantly different in Dunn’s multiple comparison test (p<0.0001). **C.** 3D segmentation of the hypocotyl epidermal cells of fixed five-day-old seedlings stained with SR2200. Cells are colored according to their volume. **D.** Quantification of cell volume and anisotropy. Graphs are combined boxplots and violin plots, n= 401-630 cells from 4-5 seedlings. Individual scatter points show outliers. Conditions not sharing a letter are significantly different according to one-way Anova with Tukeys post-hoc HSD test (p < 0.05).

### RALF perception regulates hypocotyl epidermal cell morphology and BR signaling

We sought to further investigate the molecular basis for FER function in regulating cell anisotropy. FER has been shown to bind to RALF peptides and pectin (Haruta et al. 2014; Feng et al. 2018; Xiao et al. 2019; Lin et al. 2021; Tang et al. 2021). The FER ectodomain contains two malectin-like domains, MalA and MalB (Boisson-Dernier et al. 2011). FER MalA domain was shown to bind pectin *in vitro* (Feng et al. 2018; Lin et al. 2021; Tang et al. 2021), and is required for FER function in regulating pavement cell shape and root hair growth (Gronnier et al. 2022). On the other hand, FER MalB is sufficient for RALF23 responsiveness (Xiao et al. 2019; Gronnier et al. 2022). To test whether FER function in regulating cell anisotropy and BR signaling involves pectin sensing and/or RALF perception, we analyzed *fer-4* mutant complemented with a truncated version of FER lacking its MalA domain (FER^ΔMalA^) (Gronnier et al. 2022). We noticed that FER^ΔMalA^ complements BL hypersensitivity, hypocotyl cell volume and anisotropy defects observed in *fer-4* to a level that is comparable to full length FER complementation line (Figure S4, Figure 1). These observations suggest that FER function in regulating hypocotyl cell morphology and BR signaling is primarily achieved by RALF perception and that MalA-mediated pectin sensing has a minor role in this context. To confirm these observations, we analyzed *llg1-2*, a loss of function mutant of LORELEI-LIKE GPI-ANCHORED 1 (*LLG1*) which encodes for the main RALF co-receptor in vegetative tissues (Li et al. 2015; Xiao et al. 2019; Noble et al. 2022). We observed that loss of *LLG1* leads to defects in hypocotyl epidermal cell morphology that are reminiscent of *fer-4* (Figure S5 A-B). Further, similar to *fer-4* we observed that *llg1-2* is hypersensitive to exogenous BL treatment (Figure S5D) indicating that BR signaling is hyperactive in *llg1-2*. In good agreement, we observed a decrease in *CPD* and *DWF4* transcript levels in *llg1-2* (Figure S5E). Altogether, these observations indicate that hypocotyl epidermal cell morphology and BR signaling are regulated by RALF signaling.

### RALF23 promotes BRI1-BAK1 complex formation and modulates pectin methylation status

Our observations suggest that the perception of RALF peptide(s) regulates hypocotyl cell morphology and BR signaling. RALF peptides constitute a multigenic family with 37 members in Arabidopsis Col-0 (Abarca et al. 2021). Emerging evidence shows that RALFs play structural and signaling roles at the cell surface (Moussu et al. 2023; Schoenaers et al. 2023). Positively charged RALFs associate with de-esterified pectin and mutations in RALF4, RALF22 or RALF1 basic residues affects their function (Moussu et al. 2023; Schoenaers et al. 2023; Liu et al. 2024; Rößling et al. 2024). In the context of tip-growing cells, RALF4-pectin and RALF22-pectin mers associate with LRX8 and LRX1 proteins, respectively (Moussu et al. 2023). In leaves, the association of RALF1 to pectin has been proposed to form condensates nucleating FER-LLG1 complex formation (Liu et al. 2024). RALF23 is a bona fide ligand of the FER-LLG1 complex (Stegmann et al. 2017; Xiao et al. 2019). In addition, RALF23 associates with LRX3, LRX4 and LRX5 *in planta*, and the loss of *LRX3, LRX4* and *LRX5* affects RALF23 responsiveness (Zhao et al. 2018; Gronnier et al. 2022) suggesting that RALF23 binds to both LRXs in the cell wall and FER-LLG1 at the plasma membrane. In good agreement, AlphaFold-Multimer predicts binding of RALF23 to LRX3, LRX4 and LRX5 (Figure S6A). Analogous to LRX8-bound RALF4, RALF23 exposes a polycationic surface compatible with pectin binding in the predicted LRX3-RALF23, LRX4-RALF23, LRX5-RALF23 and FER-LLG1-RALF23 complexes (Figure S6A-B). Mutating surface-exposed positively charged residues of RALF23, and pharmacological inhibition of PME abolished RALF23 bioactivity (Figure S6C-D), suggesting that, as for RALF1 and RALF4 (Moussu et al., 2023; Liu et al., 2024; Schoenaers et al., 2023; Rößling et al., 2024), pectin binding is required for RALF23 function. RALF23 perception inhibits ligand-induced complex formation between FLS2 and BAK1 (Stegmann et al. 2017), and modifies the plasma membrane nanoscale organization of FLS2, BAK1 (Gronnier et al. 2022) and BRI1 (Liu et al. 2024). We next asked whether RALF23 regulates BR signaling. In co-immunoprecipitation experiments we observed that RALF23 treatment promotes BRI1-BAK1 association (Figure 4A). In good agreement, we observed that plants overexpressing RALF23-GFP showed a decrease in *CPD* and *DWF4* transcripts accumulation as well as a decrease in the accumulation of phosphorylated BES1 (Figure 4B-C). Furthermore, the macroscopic analysis of hypocotyl length and microscopy observation of hypocotyl cells showed that overexpression of RALF23-GFP increases BL responsiveness which was suppressed by introducing the *bri1-301* allele (Figure S7). Finally, we observed that the inhibition of PME alleviated the effect of RALF23 on BR signaling (Figure S8). Taken together, these results indicate that RALF23 perception promotes BRI1-BAK1 complex formation and signaling. Since the increase in BR signaling is linked to defects in cell morphology in *fer-4* we asked whether the overexpression of RALF23 affects hypocotyl cell shape. The analysis of 3D segmentation of OxRALF23-GFP hypocotyl epidermal cells showed that overexpression of RALF23 does not affect cellular morphology (Figure 5A-B). We thus hypothesized that the promotion of BR signaling by RALF23 is not caused by defects in cell morphology. Conversely, we hypothesized that promotion of BR signaling by RALF23 is linked to its signaling function. We next asked whether RALF23 perception leads to modification of the cell wall. In dot-blot assays we observed that RALF23 treatment led to a FER-dependent decrease in esterified pectin without measurable modification of PME activity in wild-type plants (Figure 5C-D). Interestingly, we observed that RALF23 promotes the accumulation of pectin with a higher degree of de-esterification, an effect particularly pronounced in *fer-4* (Figure 5C), indicating that RALF23 perception modifies pectin composition in a FER-dependent and FER-independent manner. These modifications of pectin esterification status were associated with a decrease in PME activity (Figure 5D), which we presume to indicate that the increase in LM19 signal is due to recognition of partially esterified pectin by the antibody (Christiaens et al. 2011).

**Figure 4.**
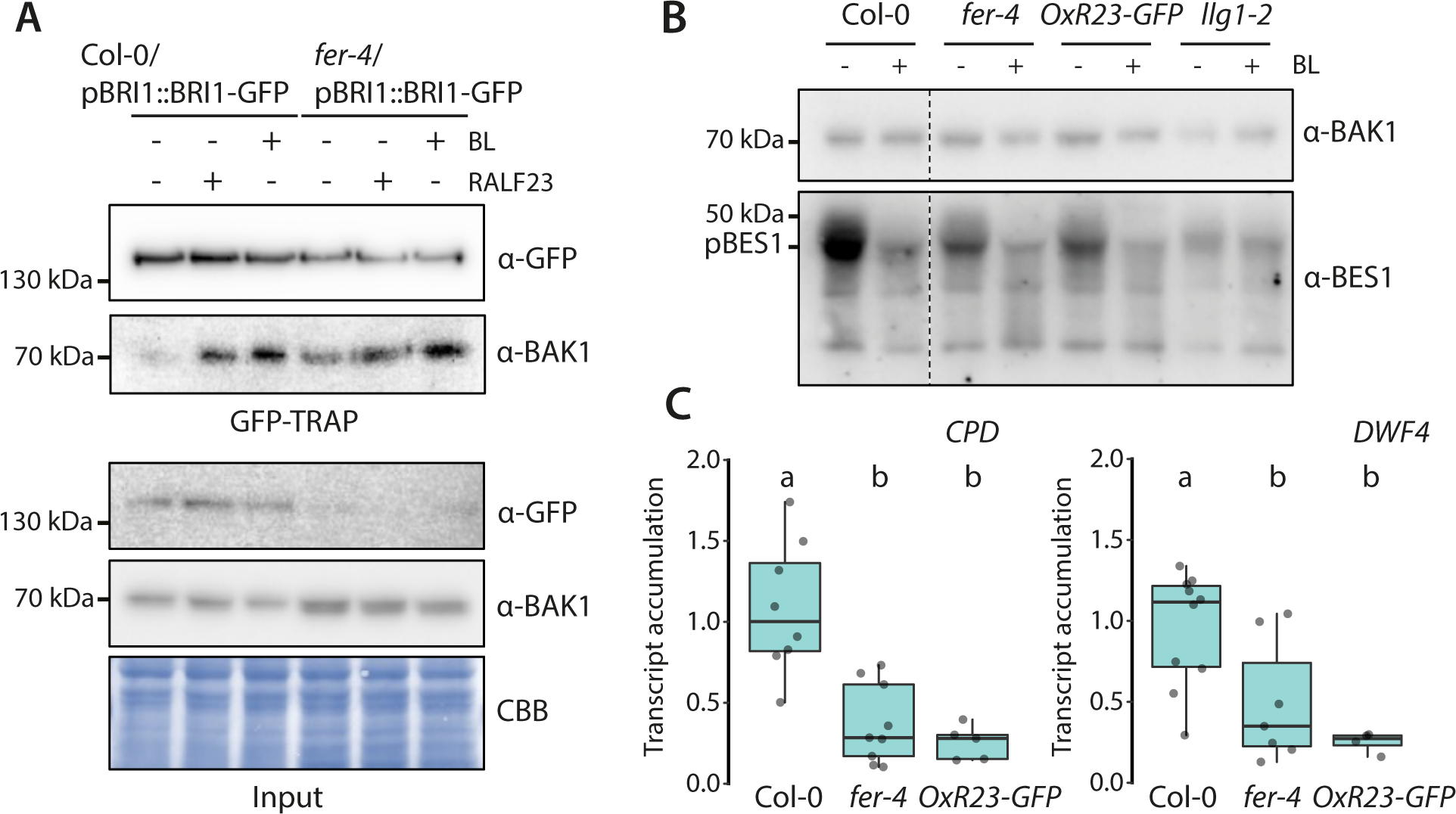
RALF23 promotes BRI1-BAK1 complex formation and signaling. **A.** Immunoprecipitation of BRI1-GFP in Arabidopsis seedlings after treatment with 1 µM BL, 1µM RALF23 or corresponding mock control solution (EtOH) for 90 minutes. Membranes were probed with anti-GFP or anti-BAK1 antibodies. Membrane stained with Coomassie brilliant blue (CBB) is presented to show equal loading. Similar observations were made in at least three independent experiments. **B.** Estimation of BES1 de-phosphorylation in twelve-day-old seedlings treated with 200 nM of BL or corresponding control EtOH for 30 min, as observed by western blot analysis using anti-BES1 antibodies. Membranes were probed with anti-BAK1 antibodies to show equal protein loading. **C.** Quantitative real-time PCR of *CPD* and *DWF4* transcripts from twelve-day-old seedlings. Scattered data points indicate measurements of individual biological samples, each corresponding to 2-3 seedlings pooled, obtained from three independent experiment. *U-BOX* was used as a house keeping gene and values are expressed relative to Col-0. Conditions not sharing a letter show significant differences according to Dunn’s multiple comparison test.

**Figure 5.**
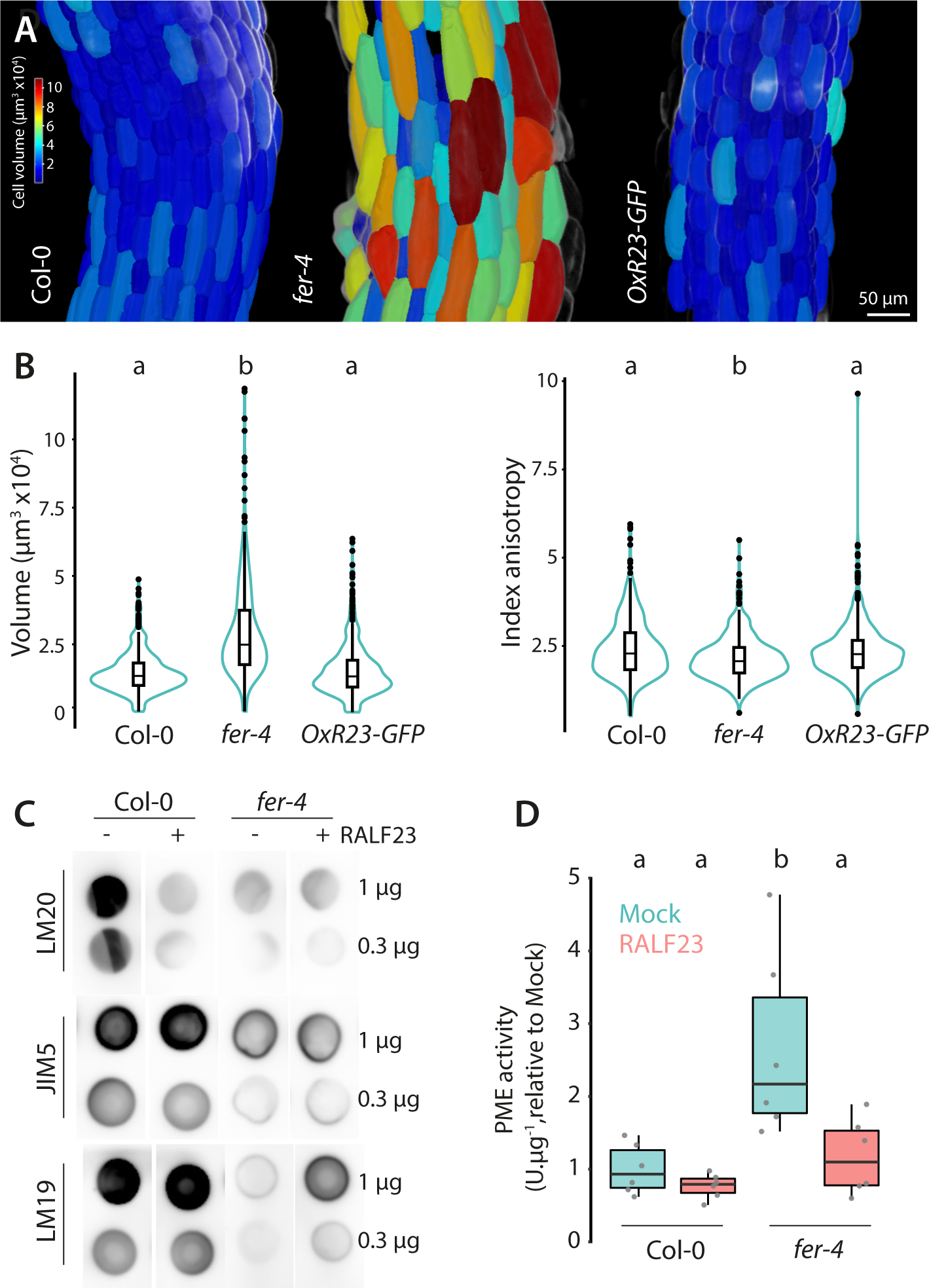
RALF23 modulates pectin methylation and the activity of pectin modifying enzymes. **A.** 3D segmentation of hypocotyl epidermis cells from fixed five-day old seedlings stained with SR2200. Heat map is colored according to cell volume. **B.** Quantification of cell volume and anisotropy. Graphs are combined box plots and violin plots, n= 377-515 cells from 3-5 seedlings. Individual scatter points show outliers. Conditions not sharing a letter are significantly different according to one-way Anova with Tukeys post-hoc HSD test (p < 0.05). **C.** Dot-Blot assays using LM20 (highly esterified pectin), JIM5 (partially de-esterified pectin) or LM19 (mostly de-esterified pectin) antibodies to probe cell wall extract of twelve-day-old seedlings treated for 90min with 1 µM of RALF23 or corresponding mock condition, µg indicates the amount of total sugars spotted on membrane. **D.** Relative PME activity of protein extracts obtained from wild-type of *fer-4* seedlings treated with RALF23 peptide for 90 min or corresponding mock control, as determined by gel diffusion assay. Scattered points show data points from 3 independent biological replicates from 2 experiments. Conditions not sharing a letter are significantly different according to Dunn’s multiple comparison test (p < 0.05).

Altogether, our observations suggest a model in which RALF23 functions as a cell wall-informed signaling cue, initiating a feedback loop solicitating BR signaling, modifying the cell wall, and coordinating cell morphogenesis (Figure 6). Our data and previous studies (Stegmann et al. 2017; Zhao et al. 2018; Xiao et al. 2019; Gronnier et al. 2022) indicates that RALF23 binds to both cell wall-located LRXs and the plasma membrane localized receptor complex FER-LLG1. As observed with other positively charged RALF peptides, RALF23 activity relies on pectin; presumably through direct binding and the formation of RALF23-pectin mers. Since RALF binding affinity to LRXs is an order of magnitude higher for LRXs than for FER-LLG1 (Xiao et al. 2019; Moussu et al. 2020), we hypothesize that the availability of wall-located epitopes defines whether RALF23 is perceived by FER-LLG1. Thereby, RALF23 perception by FER-LLG1 is informative of cell wall status and initiates adaptative responses, which involved the promotion of BRI1-BAK1 complex formation and signaling. It is conceivable that in case of perturbation of the cell wall, such as upon salt or mechanical stress, RALF peptides may be released from LRXs to be perceived by FER-LLG1. We observed that RALF23 perception modulates the activity of pectin-modifying enzymes and pectin chemistry. These modifications may generate new wall-located RALF binding epitopes and, as a consequence, potentially both modulate cell wall properties and attenuate signaling from CrRLK1Ls including FER. In this scenario, RALF23 could thus finely balance its own structural and signaling function acting within a self-regulating signaling pathway. Interestingly, we observed that modulation of pectin methylation status by RALF23 occurred in a FER-dependent and independent manner. In the case of FER-independent changes in pectin, RALF23 effects may be induced locally upon binding to LRXs, or mediated by additional plasma membrane-localized receptors, such as for instance other CrRLKL1s. Our study shows that FER regulates BRI1-BAK1 complex formation and signaling, adding to the growing list of wall monitoring cell surface receptors regulating BR signaling and providing the first example of a peptide ligand-receptor complex module in this context. Whether RALF23 promotion of BRI1-BAK1 association and signaling is a consequence of RALF23-triggered changes in pectin or whether RALF23-triggered changes in pectin are partially mediated by BRI1 signaling remains to be clarified. Interestingly, a recent report showed that BR signaling modulates FER plasma membrane localization (Chaudhary et al. 2023) indicating that RALF and BR signaling regulate each other providing a molecular circuit coordinating morphogenesis. In addition, more research is required to unravel whether RALF signaling also involves the previously identified BR-regulating wall monitoring cell-surface receptors RLP44 (Wolf et al. 2014) and RFO1/WAKL22 (Huerta et al. 2023), or whether they correspond to parallel signaling pathways converging towards BR signaling. It will also be particularly interesting to reveal whether the regulation of BRI1-BAK1 association and cell anisotropy by FER involves antagonist RALF peptides, as proposed in the context of immune signaling (Stegmann et al. 2017) and recently demonstrated in the context of reproduction (Lan et al. 2023). The perception of RALF23 modifies the plasma membrane organization of several cell surface receptor such as FLS2, BAK1 and BRI1 (Gronnier et al. 2022; Liu et al. 2024). While these observations are associated with an inhibition of FLS2-BAK1 complex formation, the functional consequences of RALF23 for other LRR-RK signaling pathways remained unknown. Here we show that RALF23 promotes BRI1-BAK1 association and BR signaling and may function as tipping point in the tradeoff between immunity and growth by rewiring cell-surface signaling. A mechanistic understanding of the molecular links between wall rheology, RALF peptides signaling and the organization of the plasma membrane organization awaits further investigation.

**Figure 6.**
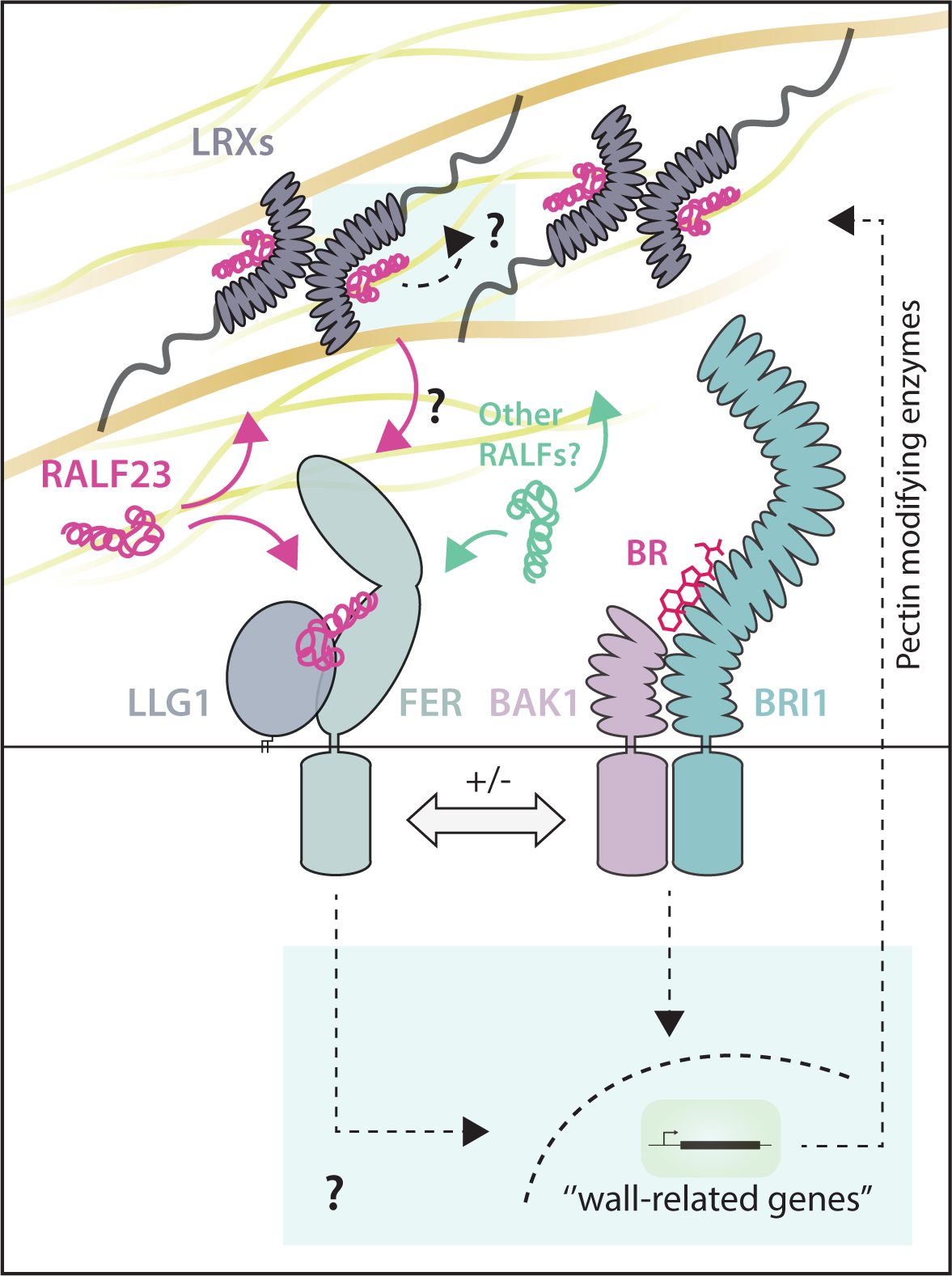
A proposed working model for a RALF-brassinosteroid morpho-signaling circuit. RALF23 binds to wall-located LRXs and/or plasma membrane localized receptor complex FER-LLG1. We hypothesize that the availability of wall-located epitopes defines whether RALF23 is perceived by FER-LLG1 or LRXs. Thereby, RALF23 perception by FER-LLG1 is informative of cell wall status and initiates adaptative responses, which involve the promotion of BRI1-BAK1 complex formation and signaling to regulate morphogenesis. RALF23 modulates pectin composition in a FER-dependent and independent manner. In the case of FER-independent changes in pectin, RALF23 effects may be induced locally upon binding to LRXs or mediated by additional plasma membrane-localized receptors such as for instance other CrRLKL1s. The definition of the exact molecular ramification connecting RALF23 signaling, BR signaling, and the cell wall status awaits further experimental investigation.

## Acknowledgements

We thank the members of the Zipfel, Grossniklaus, Ringli, Sanchez-Rodriguez, and Keller laboratories for sharing results and comments during our stimulating CCWI meetings. We thank Isabel Monte, Kyle W. Bender and Timo Engelsdorf for discussion and comments on the manuscript. We thank S. Huber for providing materials. This research was funded by the Gatsby Charitable Foundation (CZ), the University of Zürich (CZ), the University of Tübingen (JG), the European Research Council under the Grant Agreements 309858 and 773153 (grants PHOSPHinnATE and IMMUNO-PEPTALK to CZ), the European Molecular Biology Organization 2018, to JG) and the Deutsche Forschungsgemeinschaft (DFG) postdoctoral fellowship (STE 2448/1, to MS), and DFG grants (A08-SFB1101 and B01-TRR356) to JG.

## Material and methods

### Plant materials and growth

*Arabidopsis thaliana* ecotype Columbia (Col-0) was used as WT control. The *fer-4*, *fer-4*/FER-GFP (Duan et al. 2010), *fer-4*/FER^ΔMalA^-YFP (Gronnier et al. 2022), *lrx3/4/5* (Dünser et al. 2019), Col-0/p35S::RALF23-GFP (Stegmann et al. 2017), *llg1-2* (Li et al. 2015) lines were previously reported. *fer-4/bak1-4* and *fer-4/bri1-301* were obtained by crossing *fer-4* (Duan et al. 2010) with *bak1-4* (Chinchilla et al. 2007) and *bri1-301* (Xu et al. 2008) respectively. Col-0/pBRI1::BRI1-GFP (Geldner et al. 2007) was crossed *fer-4* (Duan et al. 2010) to obtained *fer-4*/pBRI1::BRI1-GFP. Seeds were surface sterilized using chlorine gas for 5h or by incubating them in 0.1 % Tween20 in 70 % EtOH for 10 min, following 70 % EtOH for 10 min and 100% EtOH for 1min. Seeds were stratified for 2 days in the dark at 4 °C and grown on half Murashige and Skoog (MS) media supplemented with vitamins, 1 % sucrose and 0.8 % agar at 22 °C and a 16-hour light photoperiod.

### RNA isolation and quantitative RT-PCR

RNA isolation, cDNA synthesis and quantitative real-time PCR (qRT-PCR) was performed as previously described (Albrecht et al. 2012b) with minor modifications. Five-day-old seedlings grown on half MS 1 % sucrose pH 5.8 were transferred to liquid half MS 1 % sucrose pH 5.8 and grown for seven days in 24 well plates. Each sample, constituted of three individual seedlings, were blot-dried, transferred to 2-mL tubes containing 2 mm glass beads and flash-frozen in liquid nitrogen. Samples were grinded while frozen for 1.5 min at 1.500rpm using BioRad TissueLyser. RNA was extracted using TRIzol™ Reagent (ThermoFischer scientific). First-strand cDNA synthesis was performed using RevertAid first strand cDNA synthesis kit (ThermoFischer scientific) and oligo(dT)18 according to the manufacturer’s instructions. cDNA was amplified in triplicate by quantitative PCR by using PowerUp™ SYBR™ Green Master Mix (ThermoFischer scientific) and a 7500 Fast real-time PCR detection system. The relative transcript accumulation values were determined by using U-box as reference and the comparative Ct method (2-ΔΔCt). The following primers were used for cDNA amplification, *CPD* (AT5G05690): forward: 5’-CCCAAACCACTTCAAAGATGCT-3’ and reverse 5’-GGGCCTGTCGTTACCGAGTT-3’, *DWF4* (AT3G50660): forward ‘5-CATAAAGCTCTTCAGTCACGA-3’ and reverse ‘5-CGTCTGTTCTTTGTTTCCTAA-3’ and *U-BOX* (AT5G15400): forward ‘5-TGCGCTGCCAGATAATACACTATT-3’ and reverse ‘5-TGCTGCCCAACATCAGGTT-3’.

### Brassinosteroid-sensitivity assay

Seeds were surface-sterilized and individually placed in line on square Petri dishes containing half MS 1% sucrose, 0.8 % phytoagar, supplemented with 200 nM epi-brassinolide or corresponding control solution (EtOH). The plates were placed at 4°C for 2 days and then placed vertically in a growth chamber for 5 days. Pictures of the plates were then taken to measure root hypocotyl length which were measured using Fiji software (Schindelin et al. 2012).

### Mobility shift-based estimation of BES1 phosphorylation status

Brassinolide-induced BES1 dephosphorylation assays were performed as previously described (Perraki et al. 2018), seedlings were germinated on half MS-agar supplemented with 1 % sucrose for five days before transplanting to 24-well plates (two seedlings per well) containing liquid half MS supplemented with 1 % sucrose. One day before the assays, the medium was replaced with fresh medium. Twelve-day-old seedlings were treated with epi-brassinolide or ethanol (mock) at the indicated concentrations for 30 min. Samples were blot-dried, transferred to 2-mL tubes containing 2 mm glass beads and flash-frozen in liquid nitrogen and store at -80 °C before protein extraction and immunoblotting.

### Co-immunoprecipitation

Co-immunoprecipitations were performed as previously described (Kadota et al. 2014). 20– 30 seedlings per plate were grown in wells of a 6-well plate for 2 weeks, transferred to 2 mM MES-KOH, pH 5.8, and incubated overnight. The next day BL (final concentration 1 µM) and/or RALF23 (final concentration 1 µM) were added and incubated for 90 min. Seedlings were then frozen in liquid nitrogen and subjected to protein extraction. To analyze BRI1-GFP/BAK1 association, proteins were isolated in 50 mM Tris-HCl pH 7.5, 150 mM NaCl, 10 % glycerol, 5 mM dithiothreitol, 1 % protease inhibitor cocktail (Sigma-Aldrich), 2 mM Na2MoO4, 2.5 mM NaF, 1.5 mM activated Na3VO4, 1 mM phenylmethanesulfonyl fluoride, and 0.5% IGEPAL. For immunoprecipitations, GFP-Trap agarose beads (ChromoTek) were used and incubated with the crude extract for 3–4 hr at 4°C. Subsequently, beads were washed three times with wash buffer (50 mM Tris-HCl pH 7.5, 150 mM NaCl, 1 mM phenylmethanesulfonyl fluoride, 0,1 % IGEPAL) before adding Laemmli sample buffer and incubating for 10 min at 95°C. Analysis was carried out by SDS-PAGE and immunoblotting.

### Immunoblotting

Protein samples were separated in 8-10% bisacrylamide gels at 150 V for approximately 2 hours and transferred into activated PVDF membranes at 100 V for 90 min. Immunoblotting was performed with antibodies diluted in blocking solution (5% fat-free milk in TBS with 0.1 % [v/v] Tween-20). Antibodies used in this study were α-BAK1 (1:5000; (Roux et al. 2011) or Agrisera AS12 1858), α-BES1 (1:1000, (Perraki et al. 2018), α-GFP-HRP (1:5000, sc-9996-HRP, Santa Cruz), α-GFP (1:5000, sc-9996, Santa Cruz). Blots were developed with Pierce ECL/ECL Femto Western Blotting Substrate (Thermo Scientific). The following secondary antibody was used: anti-rabbit IgG (whole molecule)–HRP (A0545, Sigma, dilution 1:10,000).

### Structural modelling and analysis

RALF23-LRXs and RALF23-FER-LLG1 complexes were predicted using AlphaFold Multimer (Jumper et al. 2021; Evans et al. 2022). Predicted complexes were analyzed using ChimeraX (Meng et al. 2023).

### Confocal laser scanning microscopy of propidium iodide-stained samples

Arabidopsis seedlings were stained with 50 µg/µl propidium iodide (PI) in water for 20 min, then washed with water 3 times to remove unbound PI. Samples were mounted in H_2_O on coverslips an z-stacks imaged using a Zeiss LSM880 confocal laser scanning microscope, equipped with a Plan-Apochromat 10x/0.45 M27 objective. PI was excited with a DPS 561 nm diode laser and fluorescence collected between 570-642 nm using 600-800 V gain. Z-projections of the images were created in Fiji (Schindelin et al. 2012).

### 3D Segmentation and analysis of Arabidopsis hypocotyl epidermis cells

Prior to analysis, seedlings were fixed, cleared and stained as previously described (Ursache et al. 2018) with minor modifications. Briefly, five-day-old seedlings grown on half MS-agar (0.8 % w/v) supplemented with 1 % sucrose, were fixed using 4 % Paraformaldehyde in PBS pH 6.9 while applying mild vacuum (100 mBar). Fixed samples were washed 3 times with PBS and then incubated in ClearSee solution (25 % Urea (w/v), 15 % Sodium deoxycholate (w/v), 10 % Xylitol (w/v)). Clearing was performed for at least 2 weeks while changing ClearSee every second day. Seedlings were stained using 0.1 % (v/v) SR2200 stain in ClearSee for 1 h, then washed with fresh ClearSee for at least 30 minutes. For imaging, samples were mounted in ClearSee and imaged on a Zeiss LSM880 confocal laser scanning microscope, equipped with a C-Apochromate 40x/1.2W Autocorr M27 water objective. SR2200 was excited with a 405 nm diode laser and emission detected between 413 nm-472 nm. Z-stacks acquisition was performed with an interval size of 0.3 µm to obtain cubic voxels (pixel size 0.3 µm x 0.3 µm). To analyze whole hypocotyls, z-stacks from one seedling were combined in Fiji (Schindelin et al. 2012) using the stitching plug-in (Preibisch et al. 2009). 3D segmentation and analysis were carried out in MorphoGraphX (Barbier de Reuille et al. 2015). Hypocotyl stacks were blurred using Gaussian Blur Stack with values between 1 µm-1.5 µm. Automated 3D segmentation was carried out using the Watershed Auto Seeded process with a threshold ranging from 800 – 1000. Segmentation was manually corrected, and cells not fully represented in the stack deleted. The 3D mesh was created using Marching Cubes 3D process with a cube size of 2 and 3 smoothing steps. To achieve optimal cell axis, a Bezier cord was formed according to hypocotyl shape and custom cell axis created with the process Create Bezier Grid Directions. Measurements were exported as CSV files and index of anisotropy calculated as follows: Cell Length/Cell Width = Index anisotropy.

### Pectin isolation and dot-blot assay

Pectin dot-blot assay was performed as previously described with slight modifications (Gigli-Bisceglia et al. 2022). Five-day-old Arabidopsis seedlings were transferred to liquid half MS 1 % sucrose pH 5.8 with 2 mM MES-KOH pH 5.8 and grown for six days. three seedlings per biological replicate were harvested by freezing in liquid nitrogen. Samples were incubated with 500µl protein extraction buffer (50 mM Tris-HCl pH 7.5, 150 mM NaCl, 10 % glycerol, 5 mM dithiothreitol, 1 % protease inhibitor cocktail (Sigma-Aldrich), 2 mM Na_2_MoO_4_, 2.5 mM NaF, 1.5 mM activated Na_3_VO_4_, 1 mM phenylmethanesulfonyl fluoride, and 0.5 % IGEPAL) at 8 °C for 30 min. Cell wall debris was collected by centrifugation at 16.000 g 4°C for 30min. The supernatant was collected and directly used for subsequent analysis or stored at -80 °C. Cell wall pellet was incubated in 1 ml 70 % pre-warmed (60°C) ethanol for 30 min and recovered by centrifugation at 16.000 g for 15 min. To remove residual proteins and lipids, the pellet was washed twice with 1ml Chlorform:Methanol (1:1) for 30 min. Subsequently, cell wall pellet was washed twice with 80% acetone. After removal of acetone, pellet was dried overnight at room temperature. To extract pectins, the dry pellet was boiled in 200 µl H2O for 1 h. The cell debris were collected by centrifugation (13.000 g, 15 min) and the supernatants were either directly used for western blotting or stored at -20°C. For western blotting, total sugar content was quantified using the phenol/H_2_SO_4_ method (Dubois et al. 1956) as described in (Nielsen 2010) with minor changes. 200 µl of 1:10 diluted pectin solutions were incubated with 200 µl of 5% phenol solution and 1 ml 95 % sulfuric acid. After 1 h incubation, absorbance at 490 nm was measured and total sugar content calculated using galacturonic acid as a standard. Sugar concentration was adjusted to 0.5 µg/µl and 2 µl with a total of 1µg sugars spotted on nitrocellulose membrane. Spots were dried overnight. Then, membranes were blocked with 5 % milk in PBS for 1 h. Membranes were incubated for 1 h with JIM5 (1:250 in 5 % milk in PBS), LM20 (1:250 in 5 % milk in 20 mM Tris-HCL pH 8.2, 0.5 mM CaCl_2_ and 150 mM NaCl) rat primary antibody or LM19 (1:5000 in 5 % milk in 20 mM Tris-HCL pH 8.2, 0.5 mM CaCl_2_ and 150 mM NaCl) rabbit primary antibody. Following washing with PBS + 0.1 % Tween20, membranes were incubated with anti-rat HRP-conjugated secondary antibody (1:6000 in 5 % milk in PBS, for JIM5 and LM20) and anit-rabbit HRP-conjugated secondary antibody (1:10000 in 5 % milk in PBS, for LM19) for 1 h. Once again washing was performed with PBS + 0.1 % Tween20. Blots were revealed with SuperSignal^TM^ West Pico PLUS Chemiluminescent Substrate (Thermo Scientific).

### Gel diffusion PME activity assay

Estimation of PME activity by gel diffusion assay was adapted from (Bethke et al. 2014) Proteins were extracted as described above and concentration determined using Bio-Rad protein assay kit according to the manufacturer’s instructions. Sample protein concentration was equally diluted to 0.33 µg/µl. 20 ml of a gel containing 1.2 % agarose and 0.1 % esterified pectin from apple (Sigma, 93854) and 12.5 mM citric acid, 50 mM Na2HPO4 pH 7 was prepared in 120 mm square plates. 3 mm holes were filled with 15µl of protein extract. Plates were incubated for 16 h at 37 °C. Afterwards, the plates were briefly washed and stained with 0.05% Ruthenium Red (Sigma, R2751) for 30 min. Residual dye was removed by washing. Plates were captured using a Cannon EOS 750D camera. Areas with higher staining intensity corresponding to de-esterified pectin were quantified using Fiji (Schindelin et al. 2012). Active PME units per µg of protein were calculated by using a standard curve generated with known amounts of commercial PME (Sigma, P5400).

### Statistical analysis

The number of independent experiments and the number of individual cells analyzed per condition and collected across these experiments are indicated in each figure legends. The statical tests used are reported in the figure legends and have been performed using R or GraphPad Prism.

### Ressources

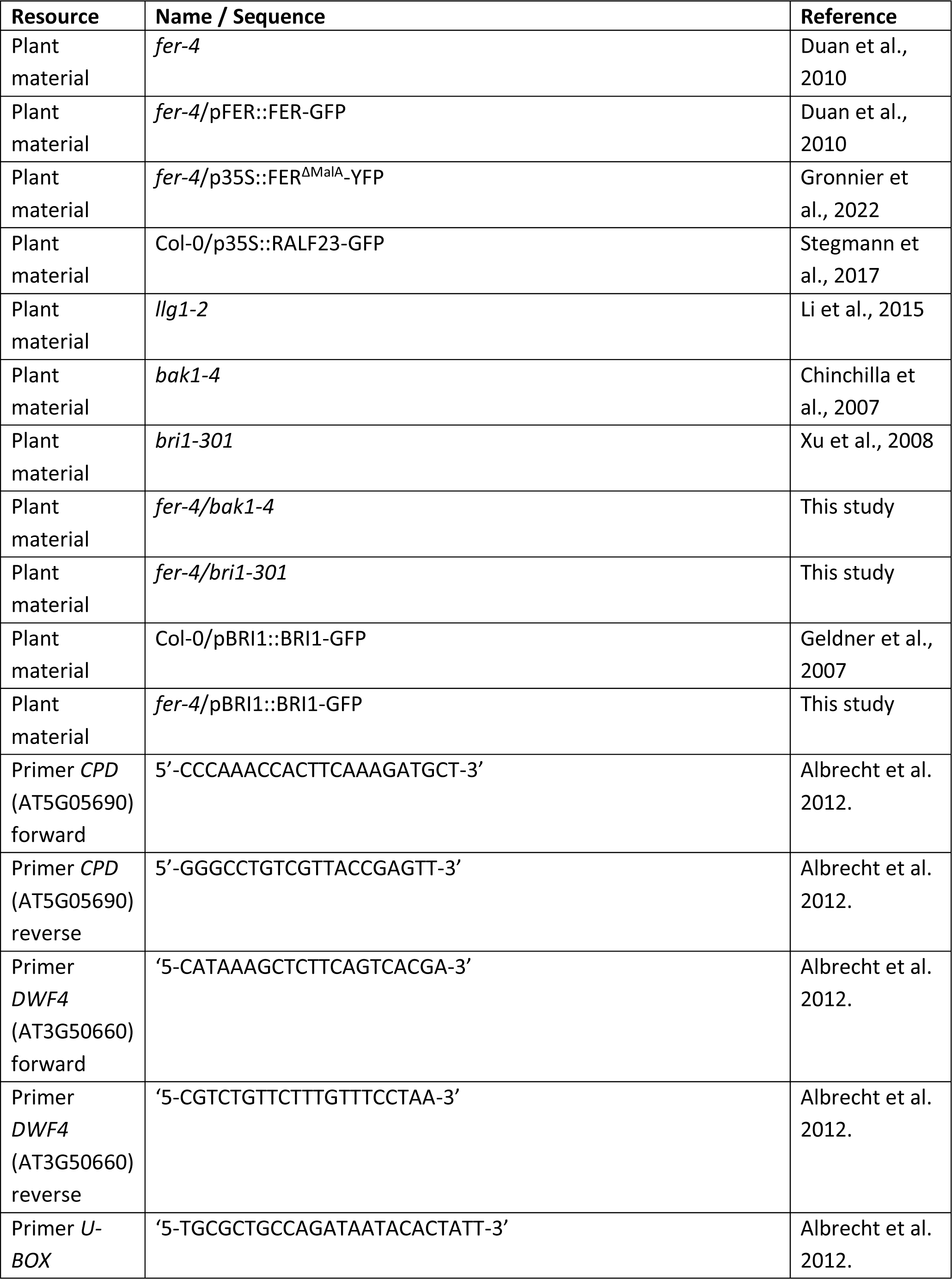
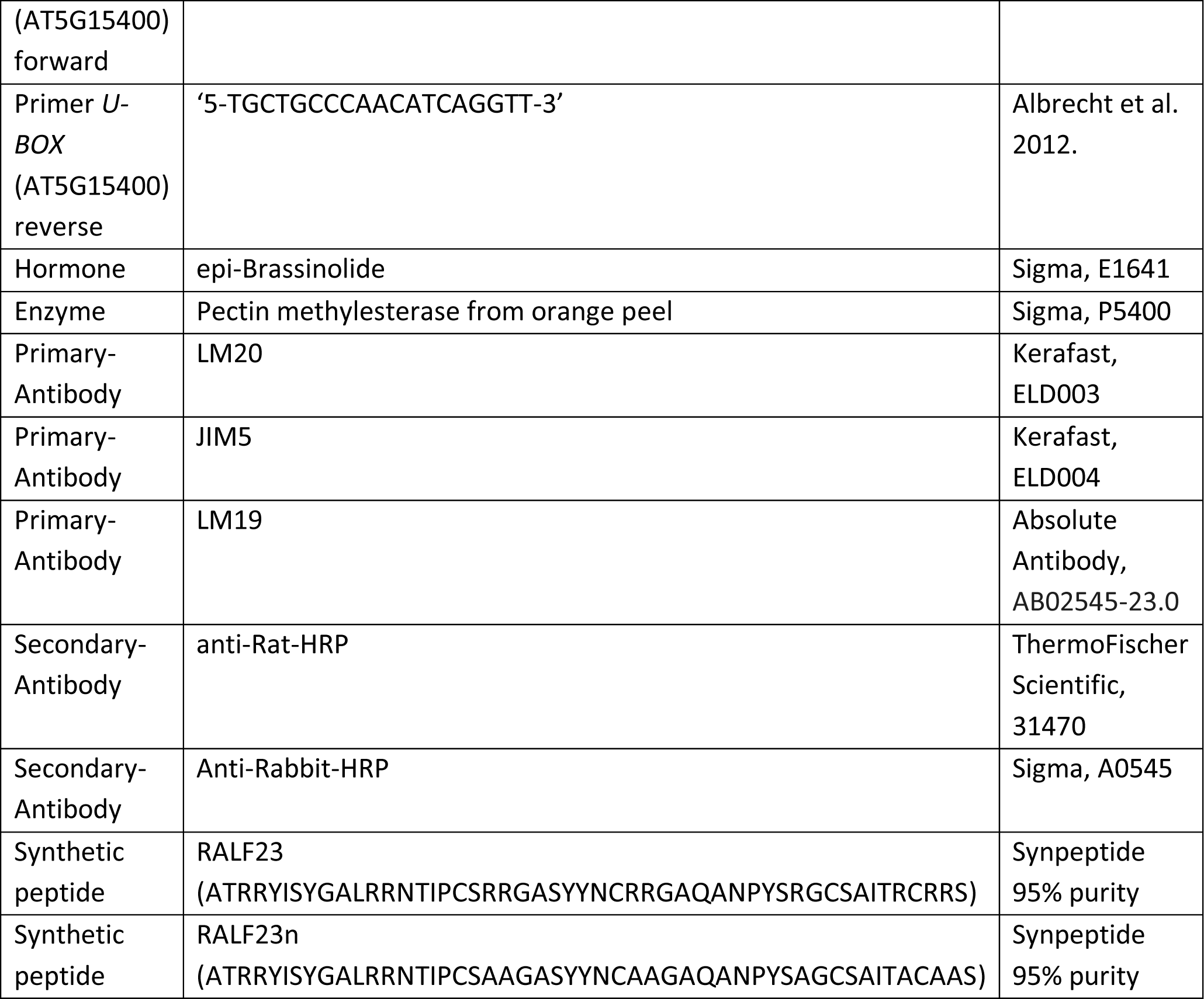

---

**Supplementary Figure 1.**
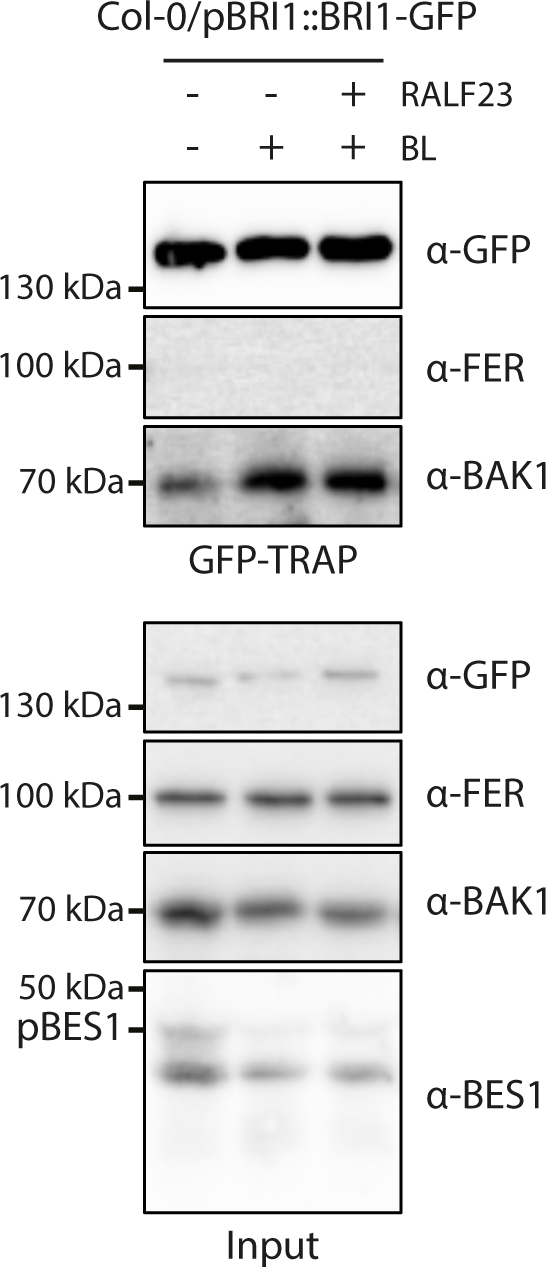
FERONIA does not associate with BRI1 receptor complex. Immunoprecipitation of BRI1-GFP Arabidopsis seedlings after treatment with 1 µM BL, 1 µM RALF23 or mock (EtOH) for 90 minutes. Blotting of membranes was performed with anti-GFP, anti-FER, anti-BAK1 or anti-BES1 antibodies. Input shows equal loading of proteins.

**Supplementary Figure 2.**
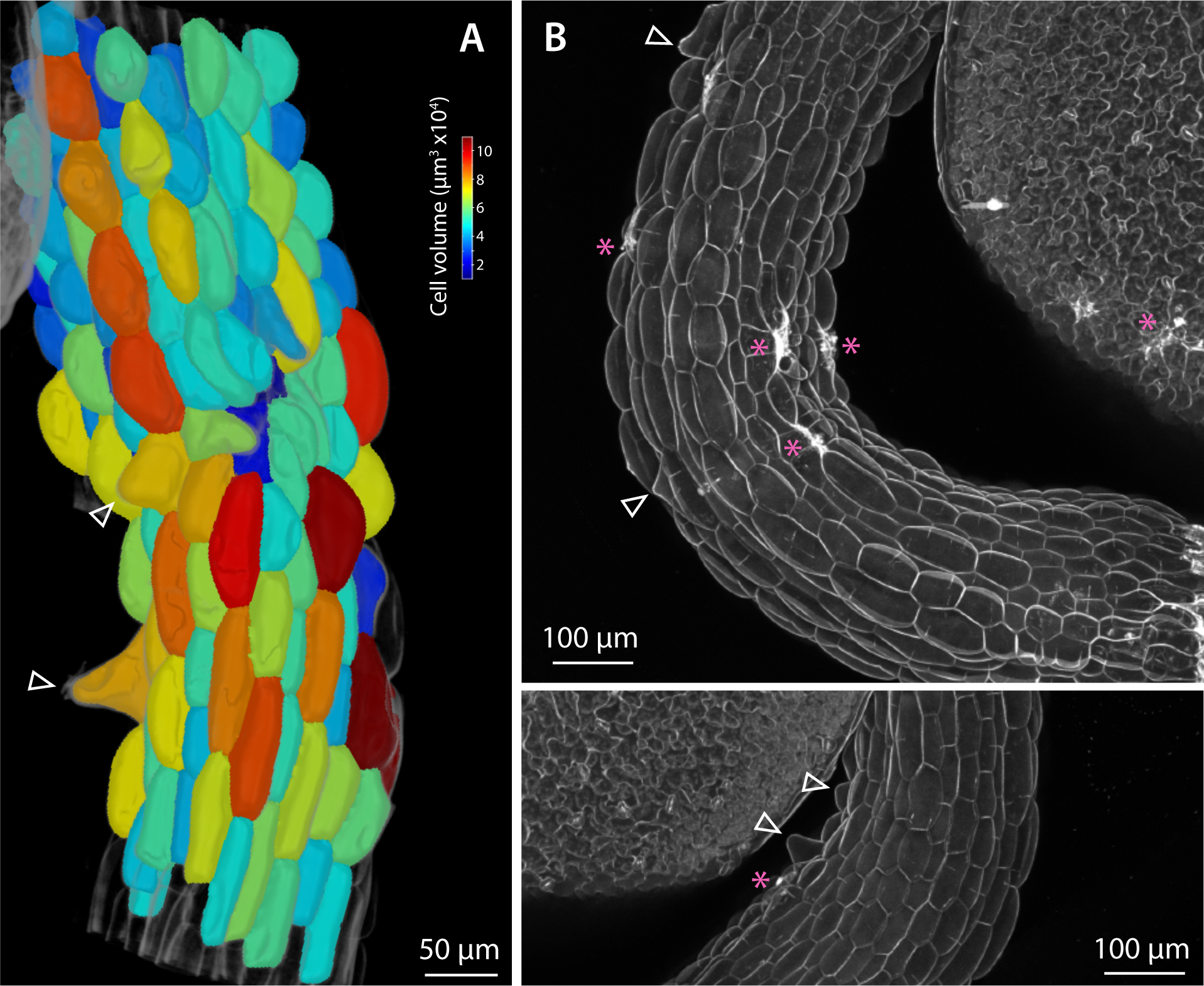
Analysis of *fer-4* hypocotyl epidermis. **A.** 3D segmentation of confocal z-stacks of SR2200-stained hypocotyl epidermis cells of fixed five-day-old *fer-4* seedlings; heat map is colored according to cell volume. **B.** Hypocotyl confocal images of five-day-old *fer-4* seedlings, stained with 50 µg/ml propidium iodide (PI). White arrows show cellular outgrowths and asterisks show intracellular PI signal which is a sign of compromised cellular integrity.

**Supplementary Figure 3.**
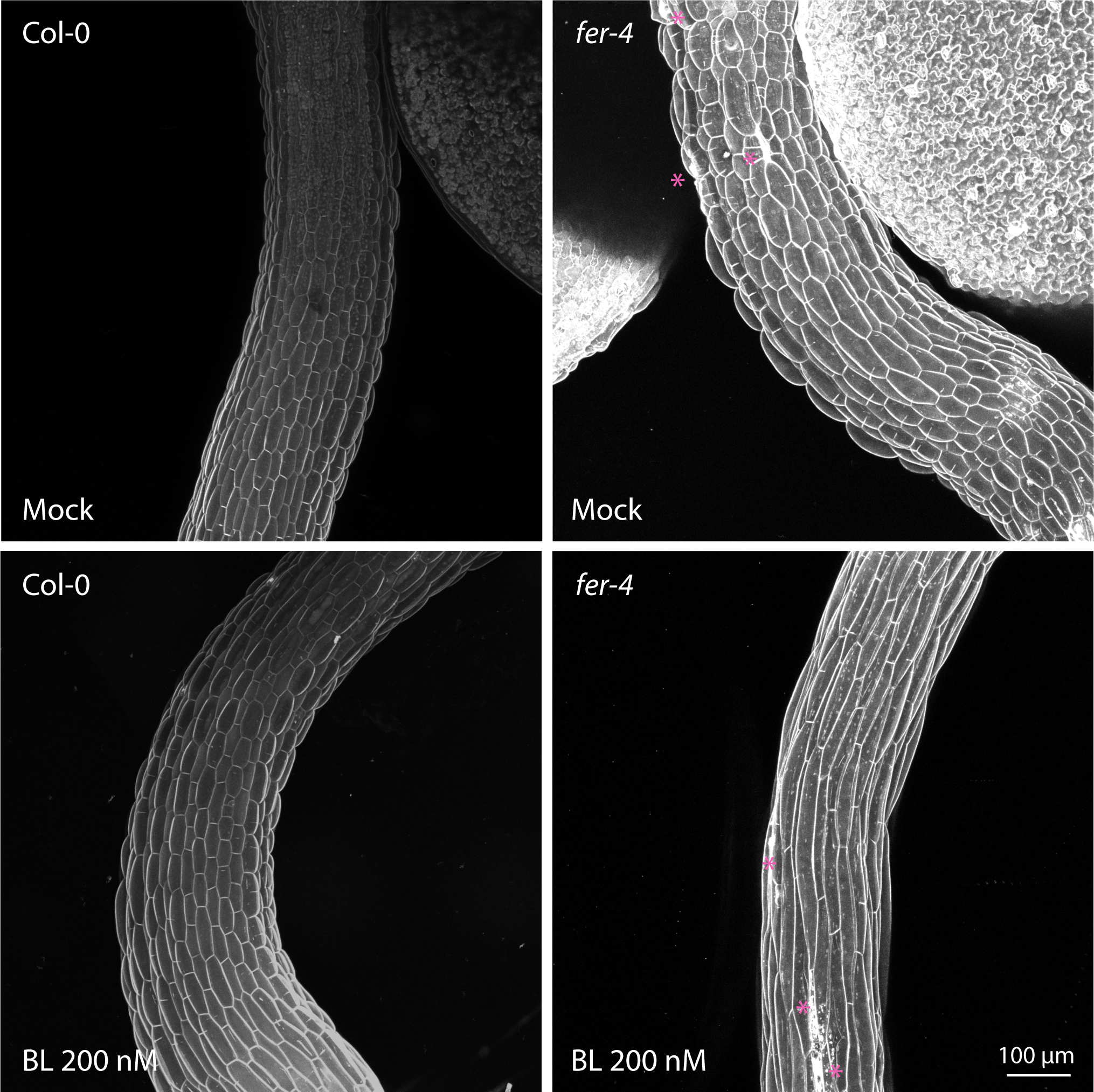
Observation of *fer-4* hypocotyl upon exogenous BL treatment. Z-projection of confocal images of three-day-old Col-0 seedlings stained with 50µg/ml propidium iodide. Seedlings were either treated with 200 nM BL or mock control solution (EtOH).

**Supplementary Figure 4.**
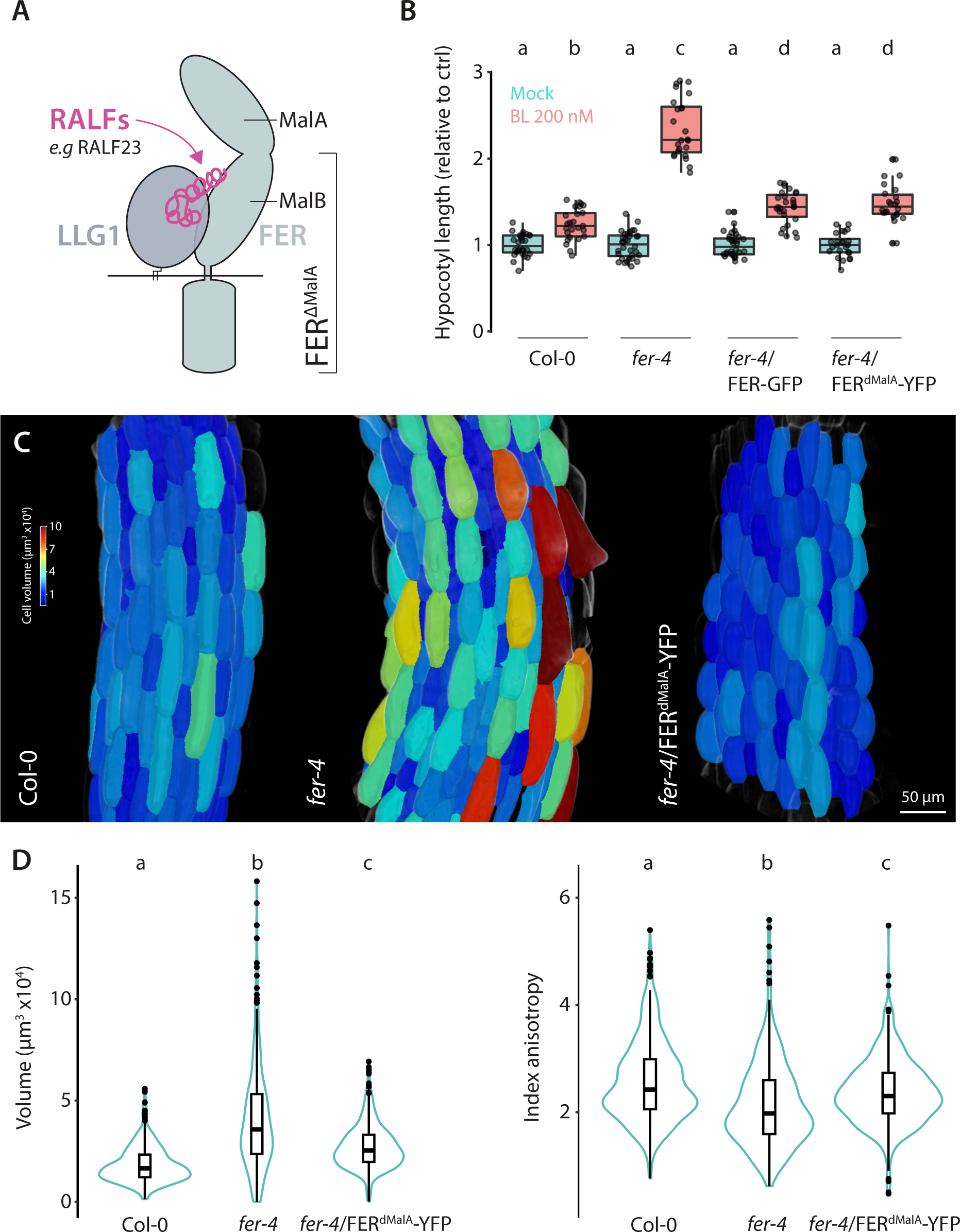
FERONIA Malectin A domain is not required to regulate cell anisotropic growth of the hypocotyl epidermal cells. **A.** Schematic representation showing FERONIA (FER) and LLG1 complex binding RALF peptides. FER^ΔMalA^ structure is indicated, the absence of MalA domain does not influence RALF binding capabilities. **B.** Quantification of hypocotyl length of five-day-old seedlings grown on half MS medium containing 200 nM BL or corresponding mock control solution (EtOH). Graphs are boxplots, scattered data points show individual measurements pooled from three independent experiments. Conditions that do not share a letter are significantly different in Dunn’s multiple comparison test (p<0.0001). **C.** 3D segmentation of hypocotyl epidermis cells of fixed five-day old seedlings stained with SR2200 and imaged by confocal microscopy. Heat map is colored according to cell volume. **D.** Quantification of cell volume and anisotropy. Graphs are violin plots, n= 277-617 cells from 4-5 seedlings. Individual scatter points show outliers. Conditions not sharing a letter are significantly different according to one-way Anova with Tukeys post-hoc HSD test (p < 0.05).

**Supplementary Figure 5.**
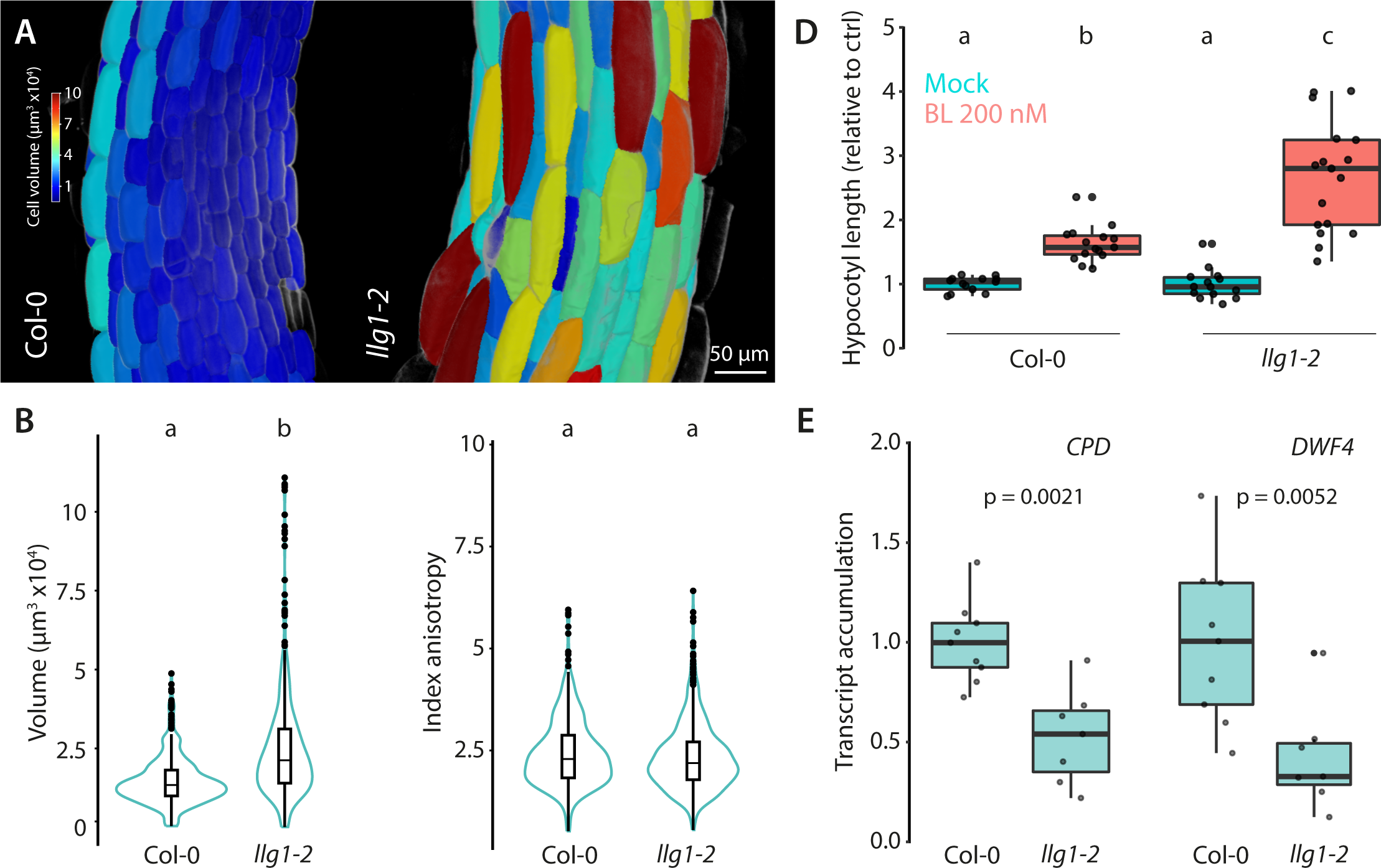
LLG1 regulates cell expansion and BRI1 signaling. **A.** 3D segmentation of hypocotyl epidermis cells of five-day old seedlings stained with SR2200 and imaged by confocal microscopy. Heat map is colored according to cell volume. **B.** Quantification of cell volume and anisotropy. Graphs are violin plots, n= 510 cells from 3 seedlings for Col-0 and 515 cells from 4 seedlings for *llg1-2*. Individual scatter points show outliers. Conditions not sharing a letter are significantly different according to one-way Anova with Tukeys post-hoc HSD test (p < 0.05). **C.** Quantification of hypocotyl length of five-day-old seedlings grown on half MS medium containing 200 nM BL or corresponding mock control solution (EtOH). Graphs are boxplots, scattered data points show individual measurements pooled from three independent experiments. Conditions that do not share a letter are significantly different in Dunn’s multiple comparison test (p<0.0001). **D.** RT-qPCR analysis of *CPD* and *DWF4* transcripts accumulation in twelve-day-old seedlings. Scattered data points indicate measurements of individual biological samples, each corresponding to 2-3 seedlings pooled, obtained from three independent experiments. *U-BOX* was used as a house keeping gene and values are expressed relative to Col-0. P values report results from Mann-Whitney non-parametric statistical test.

**Supplementary figure 6.**
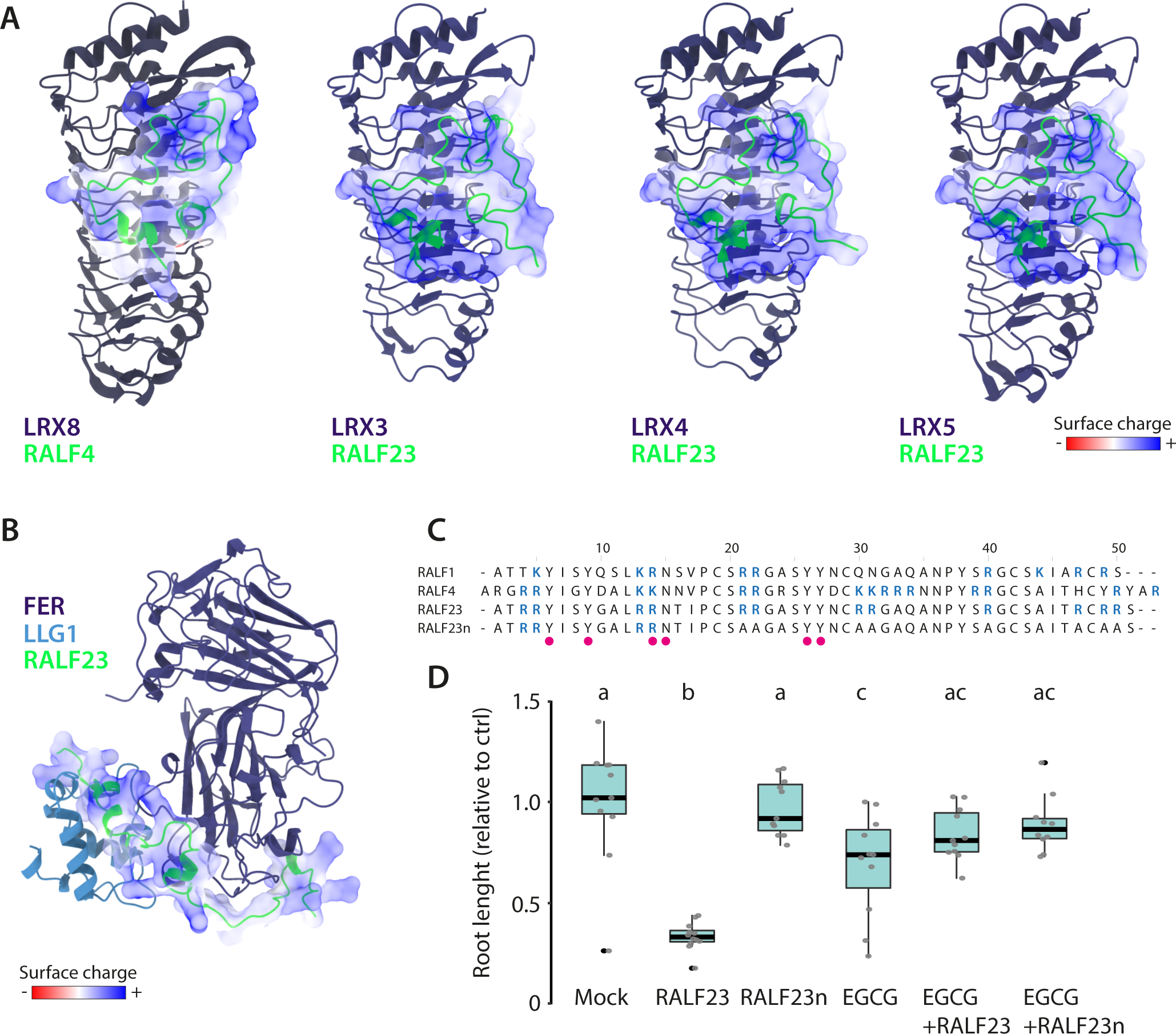
Pectin methylation status modulates RAFL23 responsiveness. **A-B**. Analysis of RALF complexes. Analysis of surface exposed residues in LRX8-RALF4 dimer 6tme (Moussu et al. 2020), and LRX3-RALF23, LRX4-RALF23 and LRX5-RALF23 (**A**) and RALF23-FER-LLG1 (**B**) predicted by AlphaFold-Multimer. RALF23 amino acids are color coded based on their charge. **C.** Sequence alignment of wild-type RALF1, RALF4, RALF23 and neutralized RALF23 (RALF23n), positively charges amino acids are indicated in blue. RALF23 residues predicted to interact with LRX3, LRX4 and LRX5 are denoted by pink circles. **D.** Relative primary root length of eight-day-old seedlings grown in the absence (mock) or presence of 1 µM RALF peptides, epigallocatechin-3-gallate (EGCG) or a combination of thereof. Graphs are boxplots, scattered data points show individual measurements pooled from two independent experiments. Conditions that do not share a letter are significantly different in Dunn’s multiple comparison test (p<0.001).

**Supplementary figure 7.**
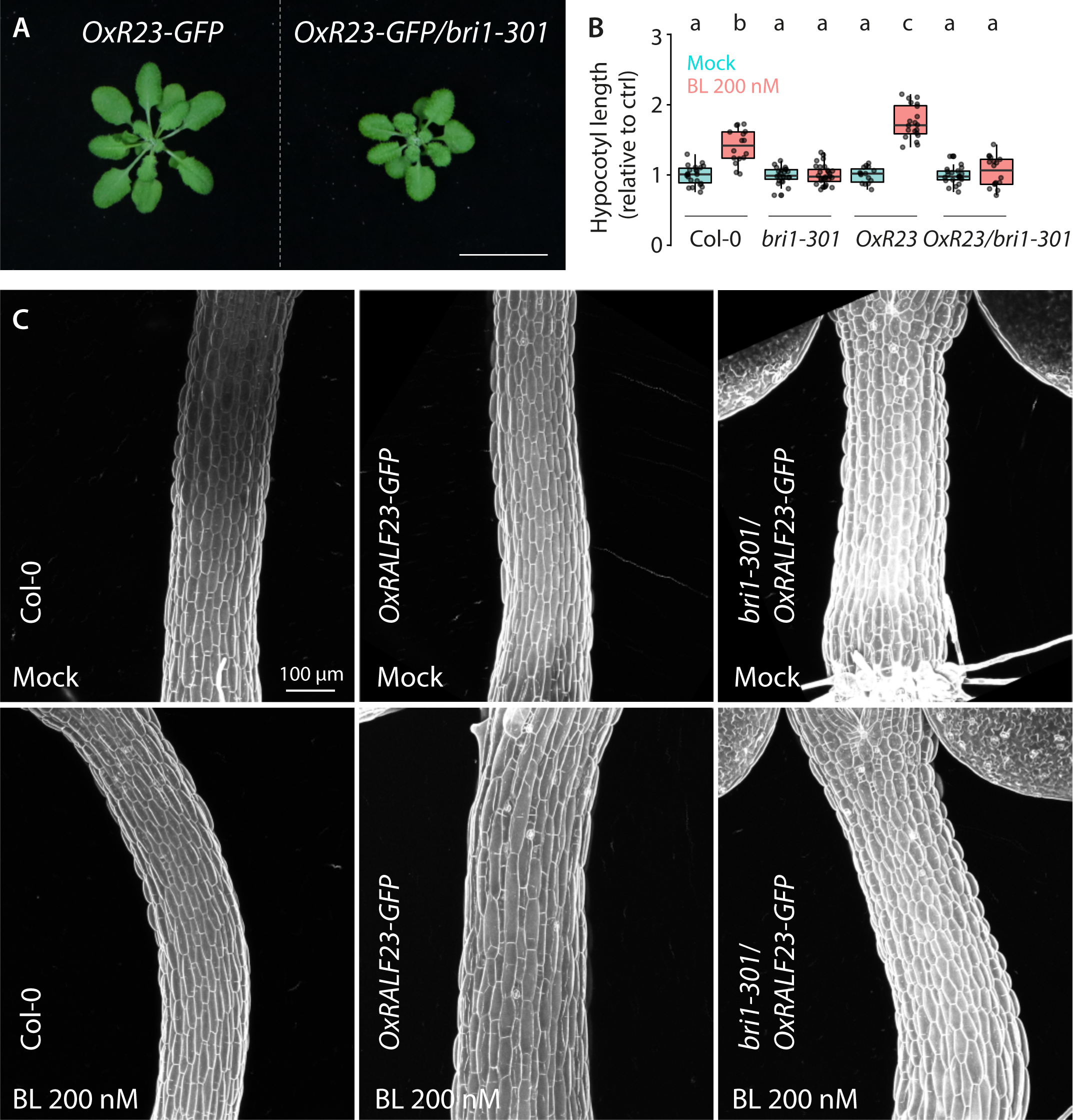
RAFL23-GFP overexpression promotes BL responsiveness. **A.** Four-week-old Arabidopsis rosettes grown in a 16-hour light daily regime. **B.** Hypocotyl length quantification of five-day old seedlings grown on half MS medium containing either 200 nM BL or mock control solution (EtOH). Graphs are presented as boxplots; scattered data points indicate individual measurements pooled from three independent experiments. Conditions not sharing a letter are significantly different in Dunn’s multiple comparison test (p<0.0001). **C.** Confocal images of three-day-old Arabidopsis hypocotyls stained with 50 µg/ml propidium iodide. Seedlings were grown on half MS medium supplemented with 200 nM BL or control solution (EtOH).

**Supplementary figure 8.**
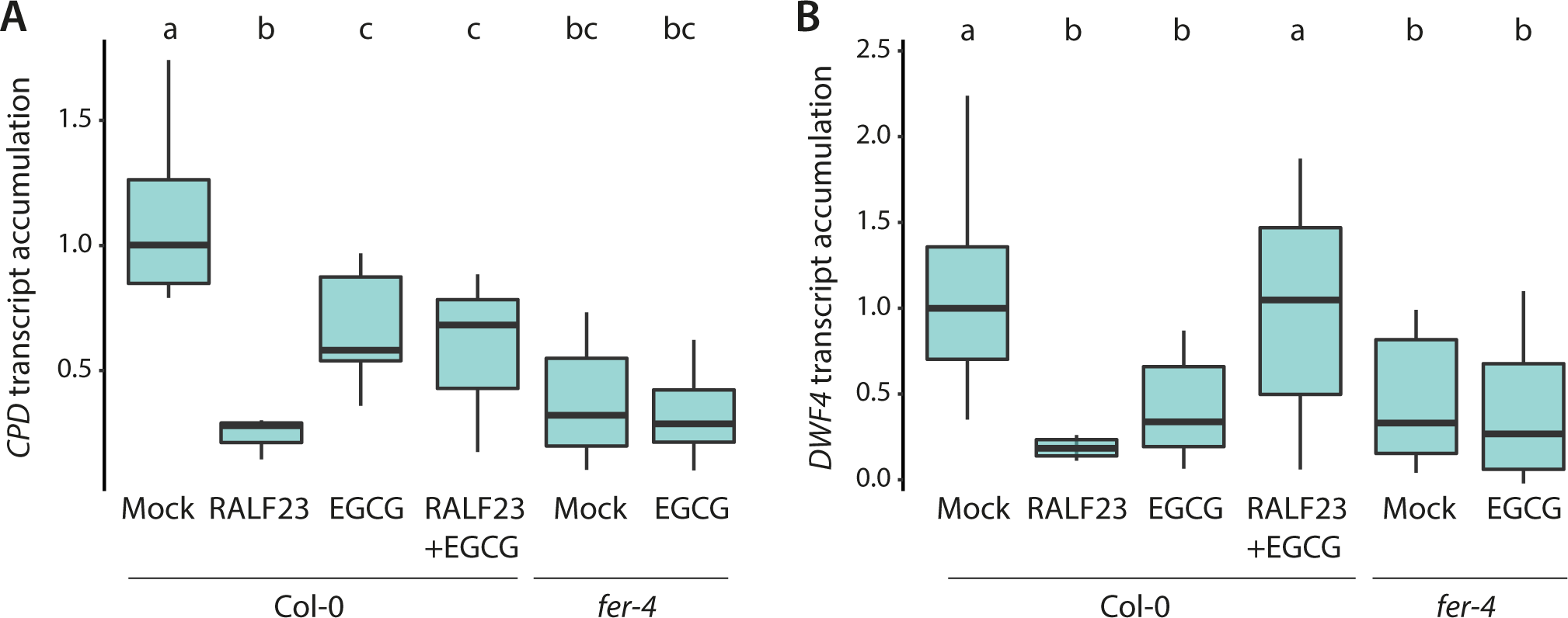
EGCG inhibits the effect of RALF23 on BR signaling. Quantitative real-time PCR of *CPD* (**A**) and *DWF4* (**B**) transcripts from twelve-day-old seedlings. Data show box plot of measurements of individual biological samples obtain from two independent experiments. Conditions which do not share a letter are significantly different in Dunn’s multiple comparison test (p<0.05).

